# Neuron cilia constrain glial regulators to microdomains around distal neurons

**DOI:** 10.1101/2023.03.18.533255

**Authors:** Sneha Ray, Pralaksha Gurung, R. Sean Manning, Alexandra Kravchuk, Aakanksha Singhvi

**Author notes:** To whom correspondence should be addressed: | |.

## Abstract

Each glia interacts with multiple neurons, but the fundamental logic of whether it interacts with all equally remains unclear. We find that a single sense-organ glia modulates different contacting neurons distinctly. To do so, it partitions regulatory cues into molecular microdomains at specific neuron contact-sites, at its delimited apical membrane. For one glial cue, K/Cl transporter KCC-3, microdomain-localization occurs through a two-step, neuron-dependent process. First, KCC-3 shuttles to glial apical membranes. Second, some contacting neuron cilia repel it, rendering it microdomain-localized around one distal neuron-ending. KCC-3 localization tracks animal aging, and while apical localization is sufficient for contacting neuron function, microdomain-restriction is required for distal neuron properties. Finally, we find the glia regulates its microdomains largely independently. Together, this uncovers that glia modulate cross-modal sensor processing by compartmentalizing regulatory cues into microdomains. Glia across species contact multiple neurons and localize disease-relevant cues like KCC-3. Thus, analogous compartmentalization may broadly drive how glia regulate information processing across neural circuits.

## INTRODUCTION

The human brain has about a 100 billion neurons and glia (von Bartheld, Bahney, & Herculano-Houzel, 2016). Glia associate closely with neurons to regulate neuron shape and function, thereby impacting animal behavior (Barres, 2008). Glia do so by molecularly modulating neuron receptive sites (NREs) through extracellular ion buffering, neurotransmitter uptake, cell-signaling, and release of neuroactive substances (Allen & Eroglu, 2017). The observed complexity in both physical and molecular glia-neuron interactions raises a fundamental organizational logic question: does a given glia regulate all contacting neurons/NREs similarly or differently?

This is relevant because across both central and peripheral nervous systems (CNS and PNS), each glia associates with multiple neurons of different sub-types. In the retina, retinal pigment epithelia glia-like cells contact NREs of rods and different cones, but whether it interacts with them similarly is unclear (Sparrow, Hicks, & Hamel, 2010). In the tongue, Type I glia-like cells contact both Type II and Type III taste cells. Finally, in the CNS, each astrocyte glia can interact with an estimate > 1000 neurons, and ~100,000 neuron receptive-endings (NREs), or sites where neurons receive information (Chung, Welsh, Barres, & Stevens, 2015). While it is known that an astrocyte glia regulates excitatory and inhibitory neurons differently (Eroglu et al., 2009; Stogsdill et al., 2017), whether or how cellular specificity dictates glia-neuron interactions at individual NREs remains largely unexplored at molecular detail.

The cellular and molecular asymmetry within both CNS and PNS glia hints that these cells can have non-uniform interactions with contacting cells. For one, glia in both settings exhibit signatures of polarity, with apical and basolateral proteins segregating into discrete glial membranes. For example, in PNS Schwann cells, apical membranes contact axons and basolateral regions face the extracellular matrix (Belin, Zuloaga, & Poitelon, 2017). Similarly, for RPE glia-like cells in the retina, basolateral proteins, such as the bestrophin chloride channel, abut the vascular choroid, while apical proteins, such as the Na+/K+-ATPase, face the photoreceptors (Gallemore, Hughes, & Miller, 1997; Strauss, 2005). CNS glia also have striking finer-grained molecular asymmetry, and localize specific molecules not only by apical-basal polarity, but into even more discrete sub-compartments, or microdomains. Thus, CNS astrocytes enrich basolateral markers such as AQP4/Aquaporins at their end-feet around epi/endothelia, Ezrin and mGluR3/5 at perisynaptic astrocytic processes, and GLT-1 and other transporters at synapses (Murphy-royal et al 2015). Drosophila ensheathing glia contain basolateral membranes, enriched with PIP3 and the Na+/K+-ATPase Nervana2, that face the extracellular matrix and an apical membrane, rich with PIP2 and sub-membranous ß_H_-spectrin, that face the neuropil (Pogodalla et al., 2021). Finally, CNS astrocytes have not only molecular, but also functional microdomains, wherein they exhibit different intracellular Ca2+ dynamics to different circuit activities (Khakh & Sofroniew, 2015). This suggests that glia differentiate inputs across contacting neurons.

*C. elegans* presents a powerful and genetically tractable experimental model to investigate specificity in glia-neuron interactions, at single-cell resolution (Oikonomou & Shaham, 2011; Singhvi & Shaham, 2019; Singhvi, Shaham, & Rapti, 2023). The animal’s nervous system comprises 56 glia and 300 neurons, each of which makes stereotypic and invariant glia:neuron contacts (Oikonomou & Shaham, 2011; Singhvi & Shaham, 2019; Singhvi et al., 2023). Furthermore, prior work has demonstrated that *C. elegans* glia regulate their associate NRE shape and function, with consequences on sensory behavior (Martin, Bent, & Singhvi, 2022; Raiders et al., 2021; Singhvi et al., 2016; Wallace et al., 2016). They also exhibit functional Ca^2+^ dynamics to a subset of neuron functions (Ding et al., 2015; Duan et al., 2020). Finally, it has been shown that loss of individual glia has varying impact on different associated neuron properties (Bacaj, Tevlin, Lu, & Shaham, 2009).

To examine specificity in glia-neuron interactions, we focused on one *C. elegans* glia cell, the amphid sheath (AMsh) glia. It is one of two glial cells in the animal’s major sense organ called the amphid, at the anterior nose-tip (Singhvi & Shaham, 2019). The AMsh glia interacts with NREs of 12 different sensory neurons, each of which senses a distinct sensory modality (Figure 1A) (Singhvi & Shaham, 2019). Thus, impact of AMsh glia on individual NREs and animal sensory behaviors can be examined reproducibly in vivo at single-cell resolution. Of the 12 neurons, AMsh glia create an autotypic channel around 8 of the NREs (Perens & Shaham, 2005; Perkins, Hedgecock, Thomson, & Culotti, 1986). Prior work has identified molecular pathways that regulate the size of the channel lumen through DAF-6/Patched and LIT-1/Nemo kinase pathways (Oikonomou et al., 2011; Perens & Shaham, 2005). Independently, we and others previously showed that AMsh glia uses the potassium chloride cotransporter KCC-3 to regulate the NRE shape and function of the animals major thermosensory neuron, the AFD, and consequently, animal thermosensory behaviors (Singhvi et al., 2016; Yoshida et al., 2016).

**Figure 1.**
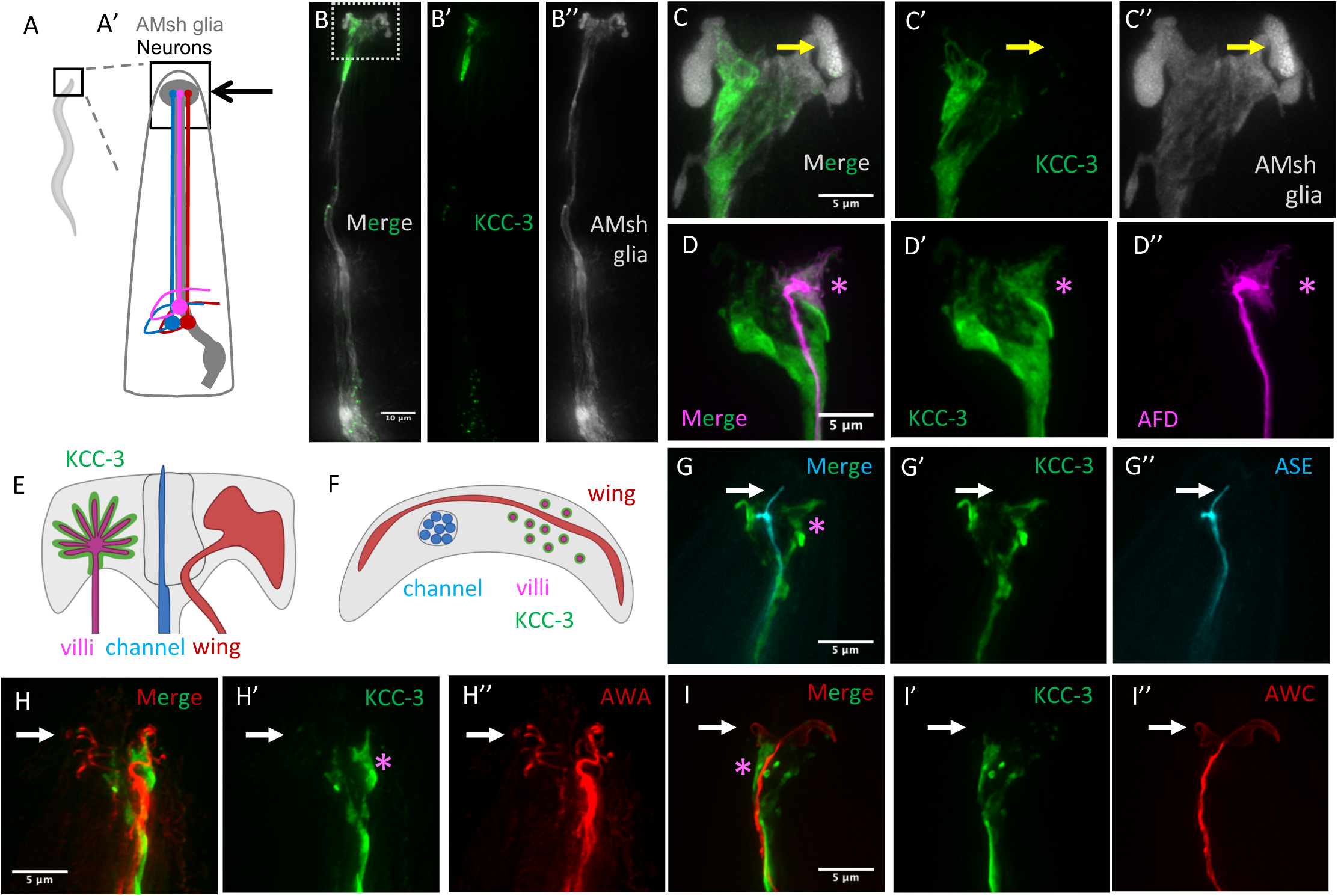
KCC-3 localizes to an apical region specifically around AFD-NRE. **(A-A’)** Schematic of a whole *C. elegans* (left) with boxed region zoomed in showing schematic of AMsh glia (grey) and three contacting neurons (magenta, red, blue). (**B-B’’**) Fluorescence image merge (B) and single channel images of KCC-3 (green) labeled by a translational mScarlet tag (B’) and AMsh glia labeled by a cytosolic CFP (B’’). Dotted boxed region is zoomed in C-I’’. Scale bar 10µm (**C-C’’**). Fluorescent image merge (C) and single-channel images of KCC-3 (green) and AMsh glia (grey) showing restricted microdomain localization. Yellow arrow notes one AMsh glial zone lacking KCC-3. (**D-D’’**) Fluorescent image merge (D) and single-channel images of KCC-3 (green, D’) and AFD-NRE (magenta, D’’) showing KCC-3 overlay on AFD-NRE region, indicated by magenta asterisk. (**E-F**) Schematic of AMsh glial contact sites with villi/AFD, channel, and embedded wing neurons as side profile (E) and top-down orthogonal view (F). Only one of the bilateral glia-neuron pair is shown. Green, KCC-3 localization schematic. (**G-I’’**) Fluorescence images as merge (G, H, I) and single-channel images of KCC-3 (green, G’, H’, I’) and NRE of ASE (cyan, G’’), AWA (red, H’’) and AWC (red, I’’) neurons. Non-overlap with KCC-3 is denoted by white arrow, site of enrichment around AFD-NRE is noted by magenta asterisk. Scale bar 5µm throughout unless otherwise noted.

Here, we report that AMsh glia compartmentalizes interactions with different NREs, with KCC-3 specifically regulating the shape of only the AFD-NRE. First, we find that KCC-3 localizes to an apical microdomain in glia. Our investigation into the molecular basis of this glia-neuron contact specificity revealed that KCC-3 localizes to a micro-domain within the glial apical membrane apposing only AFD-NRE contact sites and excluded from other NRE contact sites. Our studies find that both cell-intrinsic and non-autonomous mechanisms cooperatively drive KCC-3’s polarized apical and microdomain localization through distinct protein domains. Surprisingly, rather than AFD-NRE recruiting it, we find that cilia of heterotypic amphid NREs repel KCC-3, leading to its restrained apical micro-domain localization around AFD-NRE. Specifically, we identify the contacting ciliated NREs of two other AMsh-glia associated neurons (AWC and ASE), as sufficient to localize KCC-3 into a micro-domain. Finally, we show that this exquisitely regulated KCC-3 localization is required for its regulation of AFD-NRE shape and functions. Unexpectedly, our studies also reveal a role for KCC-3 in regulating other amphid neuron functions, including AWC, as well as interactions between the different glial regulatory cues. Taken together, these results provide molecular insight into how specificity in glia-neuron interaction is driven by compartmentalization of individual glial regulatory cues, and regulation of which impacts cross-modal sensory processing.

## RESULTS

### AMsh glial K/Cl transporter localizes to a micro-domain around the AFD neuron

We previously uncovered that the cation-chloride cotransporter KCC-3 acts in AMsh glia to regulate AFD-NRE shape and associated animal behavior (Singhvi et al., 2016). Intriguingly, we noted that a translational KCC-3:GFP reporter strain did not localize uniformly throughout the AMsh glial membrane (Figure S1A). To verify this, we engineered a dual-labeled transgenic strain with AMsh glia labeled by cytosolic CFP and KCC-3 tagged with mScarlet, and again found its localization was biased anteriorly, where the glia physically contacts 12 associated NREs, including AFD-NRE (Figure 1A-B’’).

In both confocal and 3D-SIM super-resolution microscopy, we further observed sub-cellular enrichment of KCC-3:mScarlet to a micro-domain in anterior glial membranes (Figure 1C-C’’). The pattern of KCC-3:mScarlet enrichment was anatomically reminiscent of AFD-NRE microvilli shape (Figure 1C’). We therefore performed double transgenic labeling of KCC-3 with reporters marking each of the different classes of AMsh glia-associated NREs (villi:AFD, wing:AWA/AWB/AWC, channel:ASE) (Figure 1D-I’’, S1B-B’’). This revealed that KCC-3:mScarlet predominantly localizes to glial membrane contact site only around AFD-NRE and is excluded from glial contact sites around other NREs. This localization indicates that KCC-3 is a facile molecular tool to interrogate how glia specifically target cues to subsets of contacting neurons.

### KCC-3 localizes to an apical micro-domain in AMsh

In addition to its micro-domain localization adjacent to only AFD NREs, KCC-3:mScarlet also has an anterior “tail” that extends ~50um from the animal nose tip (Figure 1B’). This tail was reminiscent of the observed expression of apical markers in glia (Low et al., 2019; Martin et al., 2022). To examine this closely, we first adapted previously reported SAX-7/L1CAM based polarity markers under the AMsh glia-specific P_F53F4.13_ promoter. As expected, we found that glial basolateral domains appose epithelia laterally and label the entire glial cell, including its cell body and process (Figure S2A). Apical membranes, on the other hand, are restricted to the anterior region of the cell, where the glia contacts NREs, and terminate in a discrete “tail” at the anterior head of the animal (Figure S2B), called hereafter the “glial apical boundary”, or GAB. Simultaneous co-labeling of both apical and basolateral domains in animals indicates that apical membranes appose neuron-contact sites while basolateral domains face outward lumen (Figure 2A-D). This localization pattern led us to posit that KCC-3 is constrained to an apical microdomain in glia.

**Figure 2:**
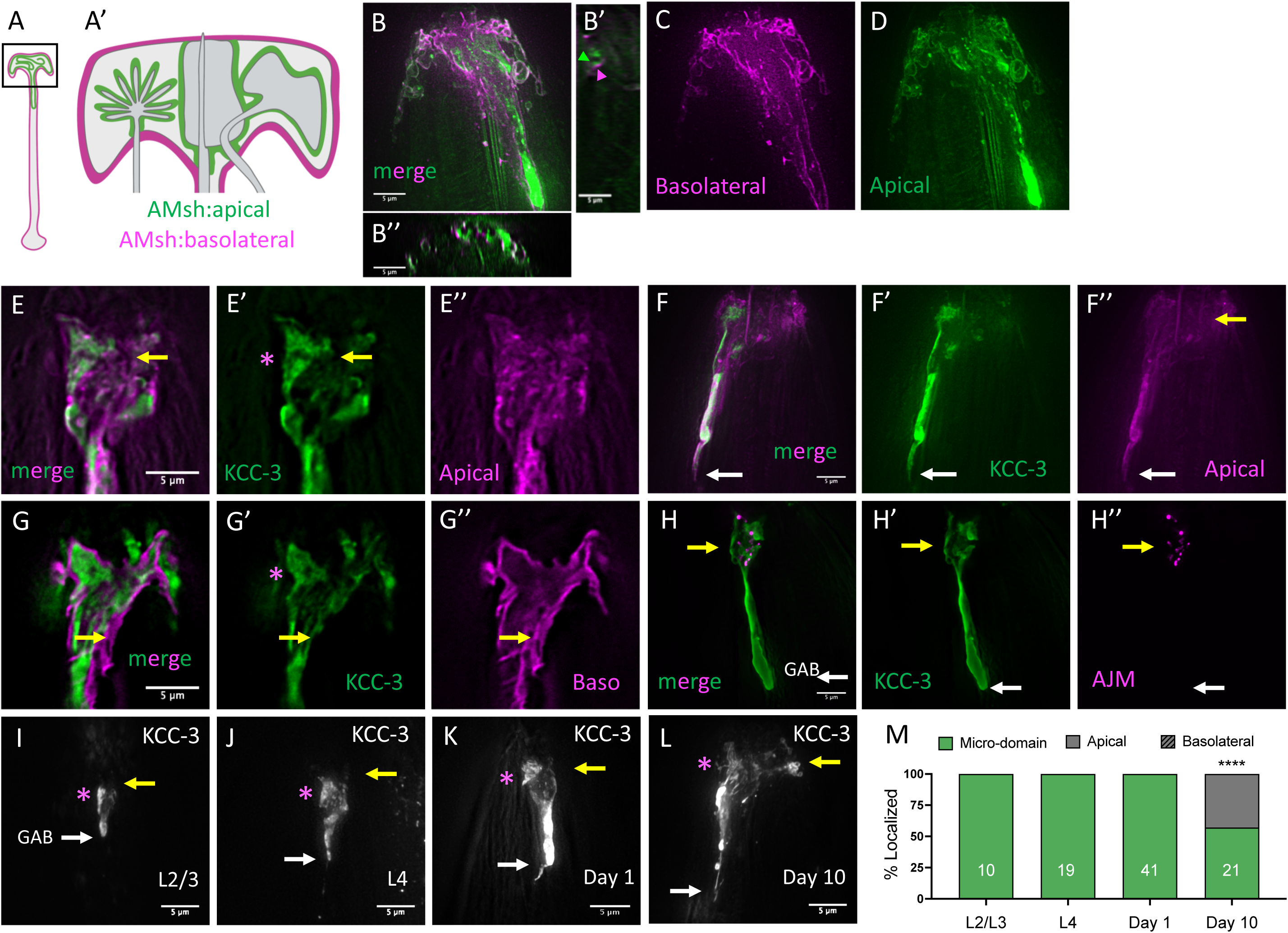
KCC-3 localizes to a glial apical microdomain in age-dependent manner. (**A**) Schematic of AMsh glia with apical and basolateral membranes marked. A= entire glia, A’= zoom of boxed region in A around anterior ending. (**B-D**) Fluorescence image overlay (B-B’’) of glial membranes marked with apical (ApiGreen) and basolateral (BasoRed) domain markers. B’ = xz orthogonal projection, B’’= yz orthogonal projection. Single-channel z-projection images of BasoRed (C) and ApiGreen (D) in AMsh glia anterior endings. (**E-F’’**) Fluorescence image overlay (E, F) of tagged KCC-3 (green E’, F’) and apical membrane marker (magenta, E’’, F’’) showing overlay with an apical microdomain. Yellow arrow denotes AMsh apical membrane lacking KCC-3. White arrow in F-F’’ denoted the GAB overlay seen in both apical marker and KCC-3 tagged reporters. Magenta asterisk in E’ and F’ denotes region of enrichment around AFD-NRE. (**G-G’**). Fluorescence image overlay (G) of tagged KCC-3 (green, G’) with basolateral membrane marker in AMsh (magenta, G’’). Yellow arrow denotes region of non-overlap. Magenta asterisk in H’ denotes region of enrichment around AFD-NRE. (**H-H’’**) Fluorescence image overlay (H) of tagged KCC-3 (green, H’) with tagged tight-junction protein, AJM-1 (H’’). Yellow arrow denotes AJM-1 staining around KCC-3 and AFD-NRE. White arrow denotes the GAB of KCC-3 past AJM-1 staining. (**I-L**). Fluorescence images of tagged KCC-3, expressed under P_AMsh_-specific promoter, showing localization as early as L2/L3 larval animals, and its aberrant expansion into non-AFD-NRE regions of the glia in aged Day10 adult animals (yellow arrow). Magenta asterisk denotes region of enrichment around AFD-NRE. White arrow denotes tail boundary. (**M**) Quantification of KCC-3 localization with age. N= number of animals on graph. **** p < 0.0001, compared to Day 1 adults. Scale bar 5µm throughout unless otherwise noted.

Since it is formally possible that SAX-7 based markers inadvertently alter cell:cell adhesion, we first chose to confirm glial apical domain identity independently. We expressed a PH-PLCδ:GFP apical membrane marker (Mahon, 2011) under an AMsh glia-specific promoter, and found that it labels the apical membrane and GAB, similar to SAX-7-based constructs (Figure S2C). Thus, this expression pattern reflects glial apical domain and is not an artefact of aberrant SAX-7 adhesion.

Next, we overlaid apical and basolateral membrane reporters with markers for AMsh-glia specific KCC-3:mScarlet. We found that the apical AMsh marker and KCC-3 colocalize at the anterior glial ending, including at the GAB, confirming that KCC-3 localizes to glial apical membranes (Fig. 2E-FG’’). Of note, however, KCC-3 expression was restricted to a sub-set of the apical membrane labeling, consistent with it being excluded from non-AFD contact sites (Fig. 2F-F’’, hash). We did not observe any KCC-3:mScarlet overlay with the basolateral AMsh membrane marker (Fig. 2H-H’’). These results show that in AMsh glia, KCC-3 localization is restricted to an “apical micro-domain” specifically at the glia’s AFD-NRE contact site.

It has been proposed that glia are analogous to neuroepithelia (Low et al., 2019). Apical-basal domains in epithelia are delimited by tight junction (Shin, Fogg, & Margolis, 2006). We therefore asked if GAB was contained by junctional proteins. However, we found that while the tight-junction marker AJM-1 localizes around the glia:NRE contact site, it does not bound the GAB domain marked by either KCC-3 or apical markers (Figure 2H-H’’). To confirm this striking result, we also examined KCC-3 expression pattern with DLG-1/DiscsLarge, another junctional marker (McMahon, Legouis, Vonesch, & Labouesse, 2001), and found again that the GAB was not delimited by DLG-1 (Figure S2D-D’’). Thus, AMsh glial KCC-3 localization to an apical microdomain is distinct from tight junction-delimited epithelia-like polarity.

We previously identified that mutations in UNC-23/BAG2 induce expansion of glial apical membranes (Martin et al., 2022). We found that loss of *unc-23* caused expansion of KCC-3 microdomains, tracking behavior of other apical markers in *unc-23* mutant animals (Figure S2E). This corroborated that KCC-3 is an apical membrane protein, and is regulated like other AMsh apical proteins at the GAB.

Finally, AFD-NRE shape deteriorates with animal age and *kcc-3* mutants exhibit age-dependent defects in AFD-NRE shape (Huang et al., 2020) We therefore wondered if KCC-3 localization tracks AFD-NRE aging. We performed longitudinal examination of KCC-3 sub-cellular localization in AMsh glia. When expressed under the heterologous AMsh-glial specific P_F53F4.13_ promoter, we observed that KCC-3 apical micro-domain in developing L2 larva that is maintained into adulthood (Figure 2I-K). As P_F53F4.13_ does not express embryonically, we determined when KCC-3 localization initates by examining rare mosaic transgenic animals with KCC-3:GFP driven under its own promoter. We found that KCC-3 expresses and localizes shortly after AMsh glia are born in the embryo (Figure S2F-G). Strikingly, the microdomain localization, but not GAB boundary, lost fidelity with age (Figure 2L-M), correlated with age-dependent decline in AFD-NRE shape (Huang et al., 2020).

### Glial KCC-3 localization is independent of AFD neuron shape or function

Given KCC-3 localization specifically around AFD-NRE, a parsimonious model would be that AFD-NRE recruits glial KCC-3 localization. Since the AFD-NRE is the primary thermosensory apparatus in the animal (Goodman & Sengupta, 2018; Kimura, Miyawaki, Matsumoto, & Mori, 2004), we first wondered if temperature regulates recruitment of glial KCC-3 to AFD-NRE contact site. However, we found that animals maintained KCC-3 apical microdomain localization, irrespective of their cultivation temperature (Figure S3A). Consistent with this, we also found that animals mutant for the sole CNG channel β-subunit required for AFD activity, TAX-2, also show intact KCC-3 localization (Coburn & Bargmann, 1996) (Figure 3A). Furthermore, KCC-3 still localizes to an apical micro-domain in *ttx-1* mutants which lack AFD-NRE microvilli that house the neuron’s sensory apparatus (Figure 3A) (Satterlee et al., 2001). Finally, the receptor guanylyl cyclase GCY-8 regulates AFD thermosensory transduction through cGMP signaling with GCY-18 and -23 (Inada et al., 2006). We previously showed that GCY-8 is inhibited directly by KCC-3-dependent chloride, and *gcy-8(ns335)* have constitutively activated cGMP production with consequently truncated AFD-NRE (Singhvi et al., 2016). Animals with a gain-of-function *gcy-8(ns335)* mutation, show intact KCC-3 localization (Figure 3A). Together, these results show that AFD-NRE shape or function do not drive KCC-3 localization.

**Figure 3:**
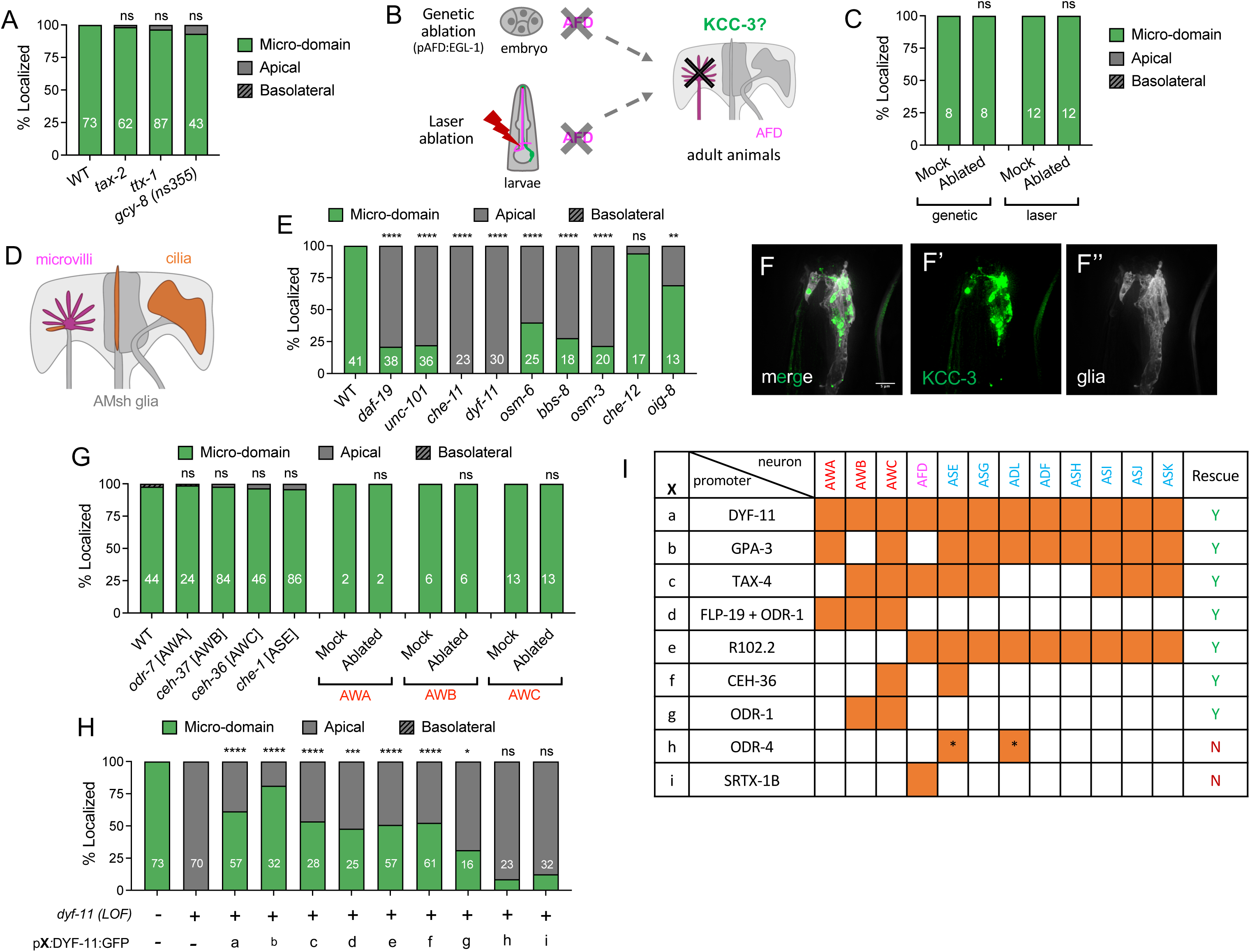
Glial KCC-3 localization is regulated by distal non-AFD-NRE cilia. **(A)** Quantification of KCC-3 localization in *tax-2, ttx-1,* and *gcy-8(ns355)* mutants compared to WT. **(B)** Schematic of genetic and laser ablation protocols to assess KCC-3 localization without AFD. **(C)** Quantification of KCC-3 localization in adults after genetic and laser ablation, compared to mock animals. **(D)** Schematic showing that all amphid NREs contain cilia. **(E)** KCC-3 localization in cilia mutants. **(F-F’’)** Fluorescence image overlay (F) of tagged KCC-3 (F’) and cytosolic glia marker (F’’) of *dyf-19* cilia mutant animals. Scale bar, 5 µm **(G)** Quantification of KCC-3 localization in amphid neuron identity mutants (*odr-7, ceh-37, ceh-36, che-1*) and after wing neuron (AWA, AWB, AWC) laser ablation. **(H-I)** Quantification of KCC-3 localization in DYF-11 rescue experiments (H). X refers to promoter(s) used for rescue experiments. Identity of X and the neurons the promoter(s) express in are expanded on in I. Orange denotes expression in associated neuron. ODR-4 expresses in an ASX and ADX neuron but the exact identity of these neurons are unclear. This is denoted by the asterisk in these boxes. For all graphs, * p < 0.05, ** p < 0.01, *** p < 0.001, **** p < 0.0001.

To test if AFD neuron altogether is dispensable, or if it has other properties driving KCC-3 localization, we next ablated the neuron altogether, in two temporally distinct ways (Figure 3B). First, we expressed the pro-apoptotic factor EGL-1 under an AFD specific-promoter (P_srtx-1_) to ablate AFD genetically (Nehme & Conradt, 2009; Singhvi et al., 2016). We first confirmed that the P_srtx-1_ promoter is expressed starting shortly after AFD’s birth in the 3-fold stage embryo, indicating that it should ablate the neuron embryonically (Figure S3B-B’’). Second, we used laser microsurgery to ablate the AFD nuclei in L1 larvae (Sulston, 1983). In either case, successful AFD ablation was tracked by complete disappearance of P_srtx-1_:GFP (Figure S3D). Surprisingly, we found that glial KCC-3 maintained restricted location to an apical microdomain in in both scenarios (Figure 3C, S3C). Taken together, our results show that while glial KCC-3 localizes to AFD-NRE contact sites, the AFD neuron or its sensory cue, NRE shape, or activity are not required for glial KCC-3 glial microdomain localization.

### Cilia of distal glia-associated neurons restrain glial KCC-3 to AFD-NRE

If AFD neuron/NRE does not recruit glial KCC-3, we hypothesized two, mutually non-exclusive, models of glial KCC-3 localization: (i) it is repelled by other NREs or (ii) its localization is regulated cell-autonomously by glia.

To test a role for other NREs, we noted that these are all derived cilia (Figure 3D). We therefore examined KCC-3 localization in animals mutant for the ciliary DAF-19/RFX transcription factor regulating all ciliary factors, and which have a complete loss of all cilia (Fig. 3E-F’’) (Perkins et al., 1986; Swoboda, Adler, & Thomas, 2000). We found that *daf-19* mutants had defects in microdomain localization, but not apical enrichment of KCC-3. Specifically, in these mutants, KCC-3 is still enriched apically, with the characteristic tail, but is no longer excluded from non-AFD-NRE contacting glial membranes (Fig. 3F-F’’).

Both ciliogenesis and transport of ciliary proteins are mediated by intraflagellar transport (IFT) bidirectionally along ciliary microtubules (Lechtreck, 2015). Briefly, IFT transport is mediated by the multi-protein subcomplexes A and B, and aided by BBS regulatory proteins, to bind cargo. Transport along microtubule tracks is guided anterogradely by Kinesin II (heterotrimer) and OSM-3 (homodimer) and retrogradely by Dynein motors. Shuttling from cell-body to cilia is guided by the clathrin coated vesicle adaptor protein-1 (AP-1) (Dwyer, Adler, Crump, L’Etoile, & Bargmann, 2001) (Figure S3E). To confirm the requirement for intact ciliary transport in glial KCC-3 localization, we performed a candidate screen of all these components. We found that loss of any of these ciliary components, including OSM-3/kinesin, CHE-11/IFT-A component, DYF-11/IFT-B component, OSM-6/IFT-B component, BBS-8/BBsome or UNC-101/AP1, led to aberrant expansion of glial KCC-3 apical microdomain localization to non-AFD-NRE contact regions (Figure 3E). Of note, except for UNC-101/AP1, enrichment around AFD-NRE was maintained (Figure S3F).

Finally, a simple caveat to these interpretations might be that the KCC-3 localization appears aberrant in cilia mutants as a secondary consequence of altered AMsh glia anterior ending shape. We therefore examined AMsh morphology in *daf-19* and *dyf-11* mutant animals. Tracking prior EM evidence (Bacaj, Lu, & Shaham, 2008; Perens & Shaham, 2005; Perkins et al., 1986)we found that AMsh glia anterior ending shape was grossly normal in both mutants, albeit marginally shrunken in *daf-19* mutant animals (Figure SG-I). Taken together, we infer that a transported ciliary protein in non-AFD NREs guides KCC-3 localization by repulsion from non-AFD-NRE contact sites, rather than recruitment to AFD-NRE.

### A two-neuron ciliary signal drives glial KCC-3 localization

We next decided to identify the neurons whose ciliary NREs regulate KCC-3. First, we examined a mutation in OIG-8/Ig domain protein, known to regulate ciliary elaboration of embedded “wing” neurons (Howell & Hobert, 2017) as well as a mutation in the CHE-12/HEAT domain protein, which only impacts “channel” neurons (Bacaj et al., 2008). *oig-8* mutants exhibit partial defects, similar to, but not phenocopying cilia mutant defects, while *che-12* mutants had no effect on KCC-3 localization (Figure 3E). A parsimonious interpretation of this data is that KCC-3 localization is driven primarily by one or more wing neurons, with channel neurons playing a lesser role. We tested this hypothesis through both candidate and unbiased cell-biology screening approaches.

First, in the candidate approach, we asked if cellular subtype identities of wing neurons are relevant. For this, we examined animals bearing mutations in ODR-7/NHR (AWA identity), CEH-37/Otx homeodomain (AWB identity), CEH-36/Otx (AWC identity) and CHE-1/GLASS Zn finger (ASE identity) (Lanjuin, VanHoven, Bargmann, Thompson, & Sengupta, 2003; Sengupta, Colbert, & Bargmann, 1994; Uchida, Nakano, Koga, & Ohshima, 2003). These genes act in parallel to DAF-19 to elaborate specific NRE cilia shapes (Figure S3J) (Lanjuin & Sengupta 2004). In these mutants, ciliary structures are altered or mis-specified into that of another “wing” neuron but are not missing. Curiously, none of these mutants perturbed KCC-3 localization (Figure 3G), suggesting that cellular identity of any of these four individual neurons is not sufficient to drive KCC-3 restricted localization. To validate this in an orthogonal approach, we also performed laser ablation of all wing neurons (AWA/B/C) individually and found that this did not alter KCC-3 localization (Figure 3G). Thus, individual wing neurons, or channel neurons alone, are not sufficient to guide KCC-3 restriction, implying that a redundant subset of neurons is required.

To identify this combination, we turned to an unbiased cell-specific rescue approach, wherein we probed which neuron expression of DYF-11 is sufficient in to rescue KCC-3 localization defect of *dyf-11* mutant animals (Fig. 3H-I). For this, we first established the validity of our approach by confirming that KCC-3 localization is rescued by DYF-11 expressed under its native P*_dyf-11_* promoter (Figure 3H-I). Similarly, DYF-11 expression under P*_gpa-3_* and P*_tax-4_* promoters, which express in 9 and 10 amphid neurons, respectively, rescues KCC-3 localization (Figure 3H-I). We next tested differing subset of amphid neurons by the non-overlapping P*_R102.2_* or (P*_flp-19_* _+_ P*_odr-1_*) combination promoters to guide DYF-11 rescuing construct in channel neurons or all wing neurons. Interestingly, both promoters rescued equally, revealing that either combination of amphid ciliary NREs can guide KCC-3. Finally, P*_ceh-36_*, which expresses in AWC+ASE, and P*_odr-1_*, which expresses in AWC+ AWB neurons, both rescued the phenotypic defects. This identifies AWC+X as a minimal neuron combination that guides KCC-3 localization. As such, expression under P*_odr-4_* (2 channel neurons) and P_SRTX-1B_ (AFD) was insufficient to rescue KCC-3 localization. These results also explain why single-neuron ablation or terminal cell-fate specification did not impact KCC-3 localization, and reveal that, minimally, AWC neurons acts with a second neuron (can redundantly be either wing or channel) to guide glial KCC-3 microdomain localization around AFD-NRE.

### Glial KCC-3 microdomain does not require canonical KCC regulators

The NRE ciliary signal needs to be received and transmitted to KCC-3 to maintain its localization. WNK and the GCK Ste20 kinases SPAK/PASK and OSR, regulate cation chloride transporters like KCC-3 across systems and species (Alessi et al., 2014; Blaesse, Airaksinen, Rivera, & Kaila, 2009; Hisamoto et al., 2008; Kaila, Price, Payne, Puskarjov, & Voipio, 2014; Payne, Rivera, Voipio, & Kaila, 2003). We therefore first asked if they are involved in this process. The *C. elegans* genome encodes a single WNK ortholog (WNK-1) (Hisamoto et al., 2008). We assessed KCC-3 localization in both a loss of function WNK-1 mutation and via RNAi and found that neither regulates AMsh glial KCC-3 localization (Figure S4A-B). Consistent with this, alignment of *C. elegans* KCC-3 with mammalian orthologs did not identify the conserved WNK and GCK kinase motif or phosphorylation sites (Figure S4C-E). We also examined mutations in the ARGK-1/creatine kinase, which localizes with KCC-3 (Burgess, Shah, Hough, & Hynynen, 2016; Salin-Cantegrel et al., 2011), and found no effect on KCC-3 localization (Fig. S4A). Finally, we have previously identified a role for the apical cytoskeletal SMA-1/β_H_-Spectrin in AMsh apical polarity regulation (Martin et al., 2022). However, animals with genetic lesions in *sma-1* exhibited normal KCC-3 localization (Figure S4A). Thus, localization of glial KCC-3 membrane transporter to apical membranes is independent of the apical SMA-1/β_H_-Spectrin cytoskeleton. Together, these results indicate that AMsh glia restricts KCC-3 localization to an apical microdomain independent of previously identified kinase regulators of cation-chloride transporters.

### A two-step model for KCC-3 apical microdomain localization

To understand how KCC-3 is localized, we decided to define the minimal KCC-3 sequences required for localization. First, we expressed *C. elegans* K/Cl homologs fluorescently tagged KCC-1 or KCC-2 under the AMsh glial promoter to ask if these sequentially similar proteins localize to an apical micro-domain like KCC-3 (Figure 4A). Both proteins in fact localized to AMsh glial basolateral membranes, in striking contrast to KCC-3 (Figure 4B-E). This indicates that sequences dissimilar between KCC-1/KCC-2 and KCC-3 drive KCC-3’s apical and micro-domain localization. Our *in silico* sequence alignment suggested that the sequence dissimilarity between KCC-2 and KCC-3 was largely restricted to three protein domains: the N-terminal, the large extracellular loop (LEL) between TM5 and TM6, and a short 81 amino acid region of the C-terminal (Figure 4F). To identify if any of these regions were relevant, we created chimera KCC proteins within each and examined localization of the chimera within AMsh glia to either a microdomain, apically, basolaterally, or elsewhere (Figure 4G-I, S6A-B).

**Figure 4:**
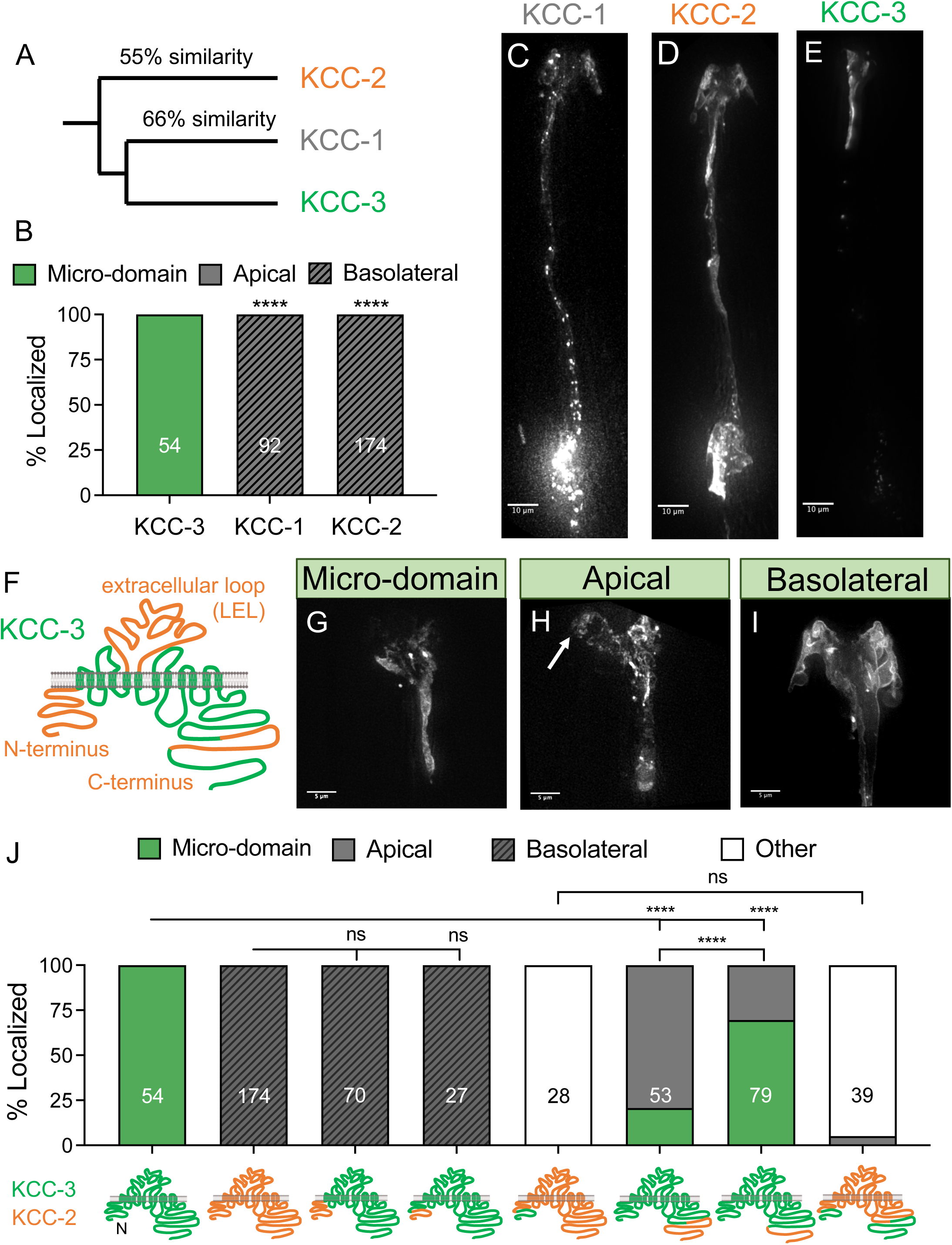
Glial KCC-3 localizes in a two-step process through two protein regions. **(A)** Phylogenetic tree denoting the relationship and sequence similarity of the three *C. elegans* KCC proteins. **(B)** Quantification of KCC-1 and KCC-2 localization when expressed in AMsh glia, compared to KCC-3. **(C-E)** Fluorescent images of KCC-1 (C), KCC-2 (D), and KCC-3 (E). **(F)** Regions of high sequence dissimilarity between KCC-3 and KCC-1/2 from *in silico* sequence alignment studies, with orange denoting regions of high sequence dissimilarity. **(G-I)** Fluorescent images of KCC localization patterns seen in KCC chimeras. White arrow points to apical expression beyond micro-domain. **(J)** Quantification of localization patterns chimeras seen in KCC chimeras. **** p < 0.0001.

First, we swapped the predicted N-terminal 90 AA sequence of KCC-3 with the 84 AA equivalent aligned sequence of KCC-2 (Figure S5A). This was sufficient to drive the chimera basolaterally (Figure 4J), suggesting that sequences contained within drive basolateral targeting of KCC-2. To narrow this further, we further swapped the first 55 amino-acid sequence of KCC-3 with the 41 amino-acid equivalent aligned sequence of KCC-2. This too drove the chimera basolaterally (Figure 4J). We divided this region further by generating a KCC-3 chimera with the first 20AA as KCC-2, but this failed to target the protein basolaterally, suggesting that the basolateral targeting sequence resides between 21-41AA in KCC-2 (Figure S6C-D). We were unable to identify a shorter basolateral targeting sequence within this 19AA KCC-2 N-terminal region by either site-directed mutagenesis of predicted phosphorylation and dileucine sites, or shorter sequence deletions (Fig. S4C-D). We also tested and found that KCC-3 N-terminal sequence is not sufficient to traffic KCC-2 protein apically (Figure 4J). Indeed, we note punctate staining in internal vesicular compartments, suggesting that lack of basolateral targeting motifs likely stall KCC-2 membrane targeting. Finally, for completeness, we also tested if it was possible that a region in KCC-3’s equivalent 55AA N-terminal sequence blocks a basolateral targeting motif. Again, we curated site-directed mutagenesis or shorter deletions in KCC-3 were unable to drive it basolaterally, suggesting this is likely not the case (Figure S6C-D). We conclude that a large and/or redundant sequence motif within a 19AA N-terminal sequence (AA21-41) drives KCC-2 basolaterally, and lack of this motif allows other domains to traffic KCC-3 apically.

To identify micro-domain motifs in KCC-3, we next engineered additional chimera proteins with varying KCC-2 C-terminal sequence. A chimera with KCC-3 sequence until the C-terminal region of high sequence dissimilarity (C-term swap A), with the last 155 AA of KCC-3 swapped with the equivalent 171 AA of KCC-2, localizes apically (Figure 4J, S5B). A chimera that with KCC-3 until just after the C-terminal region of high sequence dissimilarity (C-term swap B), with the last 68 AA of KCC-3 swapped with the equivalent 66 AA of KCC-2, exhibited faithful KCC-3 localization (Figure 4J, S5B). We therefore infer that the major microdomain-targeting motif resides in a region of high sequence dissimilarity between KCC-3 and KCC-1/2, amid amino acids 915-997 of KCC-3 (Figure S5B). However, while necessary, we note that these sequences are not sufficient to override the strong basolateral targeting sequences of KCC-2 (Figure 4J), but only operate when this basolateral motif is absent.

Since K/Cl proteins exist as oligomers (Simard et al., 2007) and the dimerization domain is thought to reside in the cytosolic C terminus, it is possible that dimerization would impact our inference of the motifs above. We asked if microdomain localization of the chimera was due to its shuttling with endogenous protein as a heterodimer. To test this, we examined localization of these chimeras in *kcc-3(ok228)* mutant animals. We found that trends hold equally in both wild type and mutant background (Figure S6E). Indeed, our results support the notion that if at all, endogenous protein may even partly hinder chimera localization. Thus, cross-oligomerization with wild-type protein cannot explain the ability of C-terminal chimeras to localize to a microdomain.

Together, these results reveal a two-step model for glial KCC-3 localization. First, lack of basolateral-targeting motifs in KCC-3 N-terminal region allows it to be shuttled apically to GAB. Once there, ciliary NRE signals act with the C-terminal 915-997 AA sequence to restrict KCC-3 to a microdomain.

### AMsh glial KCC-3 micro-domain localization regulates AFD neuron shape and function

Having investigated how, we next asked why AMsh glia and non-AFD NREs regulate KCC-3 localization. We hypothesized that it could play two roles; (a) regulation of AFD-NRE function, and (b) prevent it from aberrantly impacting other NRE functions. To test this, we examined properties of both AFD and other NREs, in *kcc-3(ok228)* mutant and mis-localized KCC-3 transgenic animals.

First, we asked if KCC-3 localization impacted AFD-NRE shape. Exploiting the fact that *kcc-3* mutants have disrupted AFD-NRE shape (Figure 5A-A’) (Singhvi et al., 2016), we examined if mis-localized KCC-3 chimeras can rescue AFD shape in a *kcc-3* mutant background. While expression of full-length KCC-3 in a *kcc-3* background rescues AFD shape, a basolaterally localized chimera (KCC-2^Nterm^ in an otherwise KCC-3 protein) fails to rescue AFD shape (Fig. 5B). In contrast, a chimera that shows apical localization was able to rescue *kcc-3(ok229)* mutant AFD-NRE defects (Figure 5B), even if slightly less efficiently than a microdomain-exclusive construct. Together, we infer that apical localization, but not necessarily micro-domain localization, KCC-3 is required for its regulation of AFD-NRE shape.

**Figure 5:**
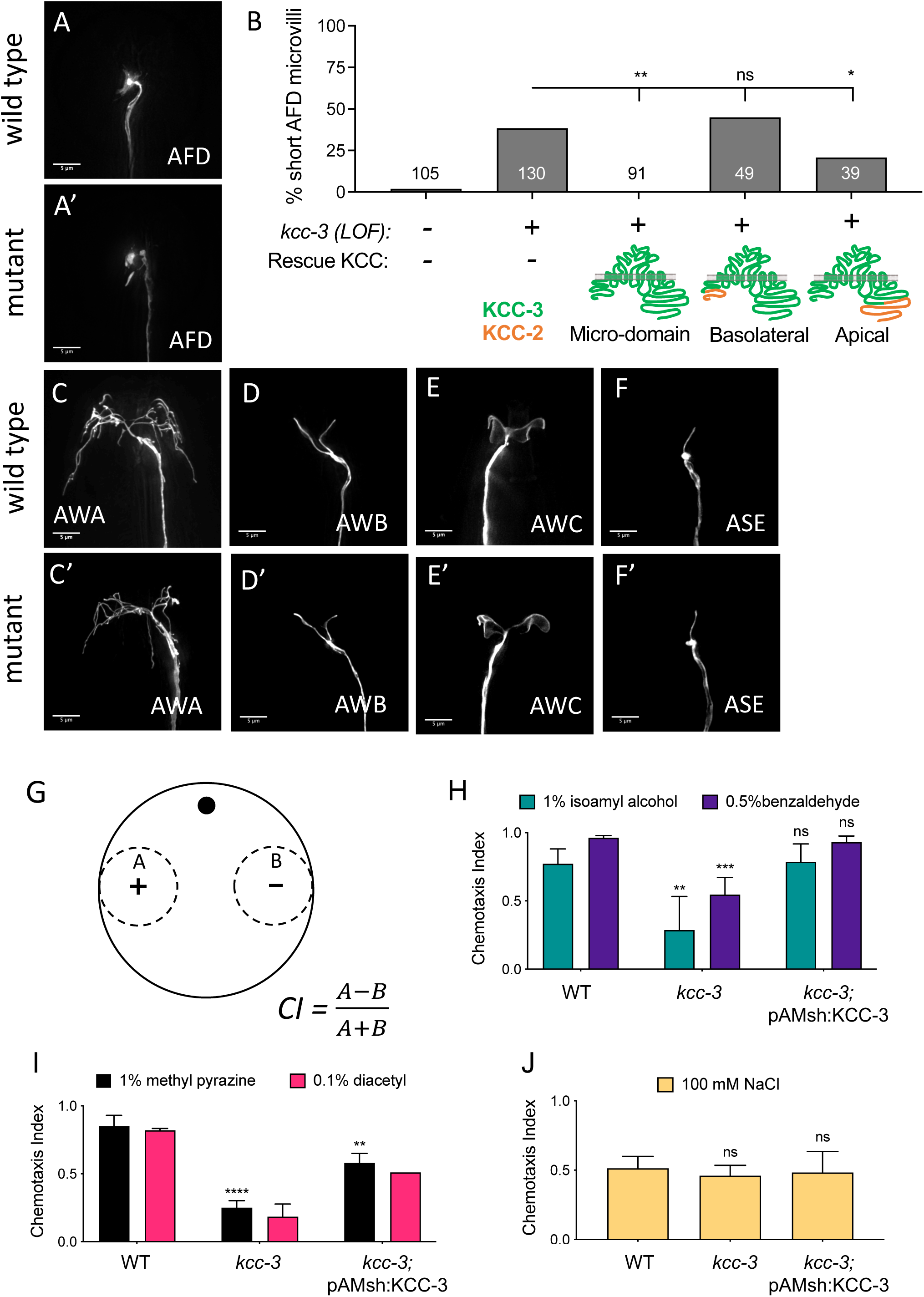
Microdomain localization of KCC-3 regulates both AFD and non-AFD neuron shape and associated animal behavior. **(A-A’)** Fluorescent images of AFD NRE in both wildtype (A) and *kcc-3(ok228)* mutants (A’). **(B)** Quantification of AFD NRE shape rescue with WT KCC-3, basolaterally localized KCC-2/KCC-3 chimera, and an apically localized KCC-2/KCC-3 chimera. **(C-F’)** Fluorescent images of non-AFD amphid NRE shape in wildtype and *kcc-3(ok228)* mutant backgrounds. **(G)** Schematic of chemotaxis assays, including equation for chemotaxis index (CI). **(H-J)** Behavioral assay quantification for AWC-sensed odorants (H), AWA-sensed odorants (I) and ASE-sensed tastant. All included at least 3 biological replicates with at least 90 animals/trial, except diacetyl, which only had 2 biological replicates. All behavioral data compared to wildtype. * p < 0.05, ** p < 0.01, **** p < 0.0001.

We also asked the corollary—is aberrant apical expansion of KCC-3 detrimental to AFD-NRE shape? *dyf-11* and *osm-6* mutant animals, where AMsh glial KCC-3 enriches around AFD-NRE besides also expanding aberrantly to other NREs (Figure 3E), do not exhibit defects in AFD-NRE shape (Singhvi et al 2016). Thus, we conclude that enrichment of KCC-3 regulates AFD-NRE shape but its expansion to other apical regions is inconsequential to the AFD.

### AMsh glial KCC-3 regulates distal AWC neuron activity

If KCC-3 expansion is inconsequential to AFD-NRE, why is it regulated? We decided to formally test if KCC-3 can impact other NRE properties, despite not localizing to their membrane contact sites (Figure 1G-I’’, S1B-B’’). As expected, we found that loss of *kcc-3* does not impact the shape of the wing neurons (AWA, AWB, AWC), and the channel neuron ASE (Figure 5C-F’, S7A). Surprisingly, however, *kcc-3(ok228)* null mutant animals exhibited behavioral defects in *kcc-3* mutants for wing neuron-driven animal behaviors. Specifically, *kcc-3* mutant animals fail to chemotax towards the AWA-sensed odorants methyl pyrazine and diacetyl and the AWC-sensed odorants isoamyl alcohol and benzaldehyde (Figure 5G-I). These deficits were comparable to defects in AMsh glia-ablated animals and previously reported behavior loss of DYF-11/IFTB (Figure S7B-C) (Bacaj et al., 2008, 2009). Animal behavior mediated by the ASE-channel neuron sensed tastant 10mM NaCl is, however, unaffected (Figure 5J).

We asked if these defects were secondary to *kcc-3(ok228)* AFD-NRE defects through indirect electrical or chemical synaptic deficits between AFD and other neurons. To parse this, we tested *ttx-1* mutants, which lack AFD-NRE for these behaviors and observed normal chemotaxis behaviors towards methyl pyrazine, isoamyl alcohol, and benzaldehyde (Figure S7B). Thus, defects in AFD-NRE shape or functions alone cannot explain the observed deficits in AWA/C functions.

Next, we wondered if the defects arose from KCC-3 requirement in AMsh glia, or its indirect function in other glia that associate with the downstream circuit interneurons (Singhvi & Shaham, 2019; White, Southgate, Thomson, & Brenner, 2008). To test this, we asked if expression of KCC-3 only in AMsh glia could rescue the AWA/C wing-neuron behavior defects. We found that it could for AWC-dependent behaviors but only partially rescues AWA- or AWB-dependent sensory animal behaviors (Figure 5H-I). We infer that AMsh glial KCC-3 can indirectly affect AWC function, despite not localizing to the glial contact site of this neuron/NRE. Defects in AWA-dependent behaviors may arise from a combined requirement of KCC-3 in AMsh and other glia.

We therefore decided to focus on AWC, and examine how its neuron activity profiles track KCC-3 by functional imaging of intracellular Ca^2+^ dynamics using a cell-specific expression of fluorescent reporter GCaMP (Chalasani et al., 2007). Tracking animal behavior data, we observed attenuated reponses of the AWC neuron to iso-amyl alcohol (Figure 6A, 6C, 6D). More interestingly, when challenged with the odor a second time with a 30s interval, the attenuation in AWC responses was larger (Figure 6B). Thus, while not present at its contact-sites, glial KCC-3 can nonetheless regulate neuron response properties of the distal AWC-NRE.

**Figure 6:**
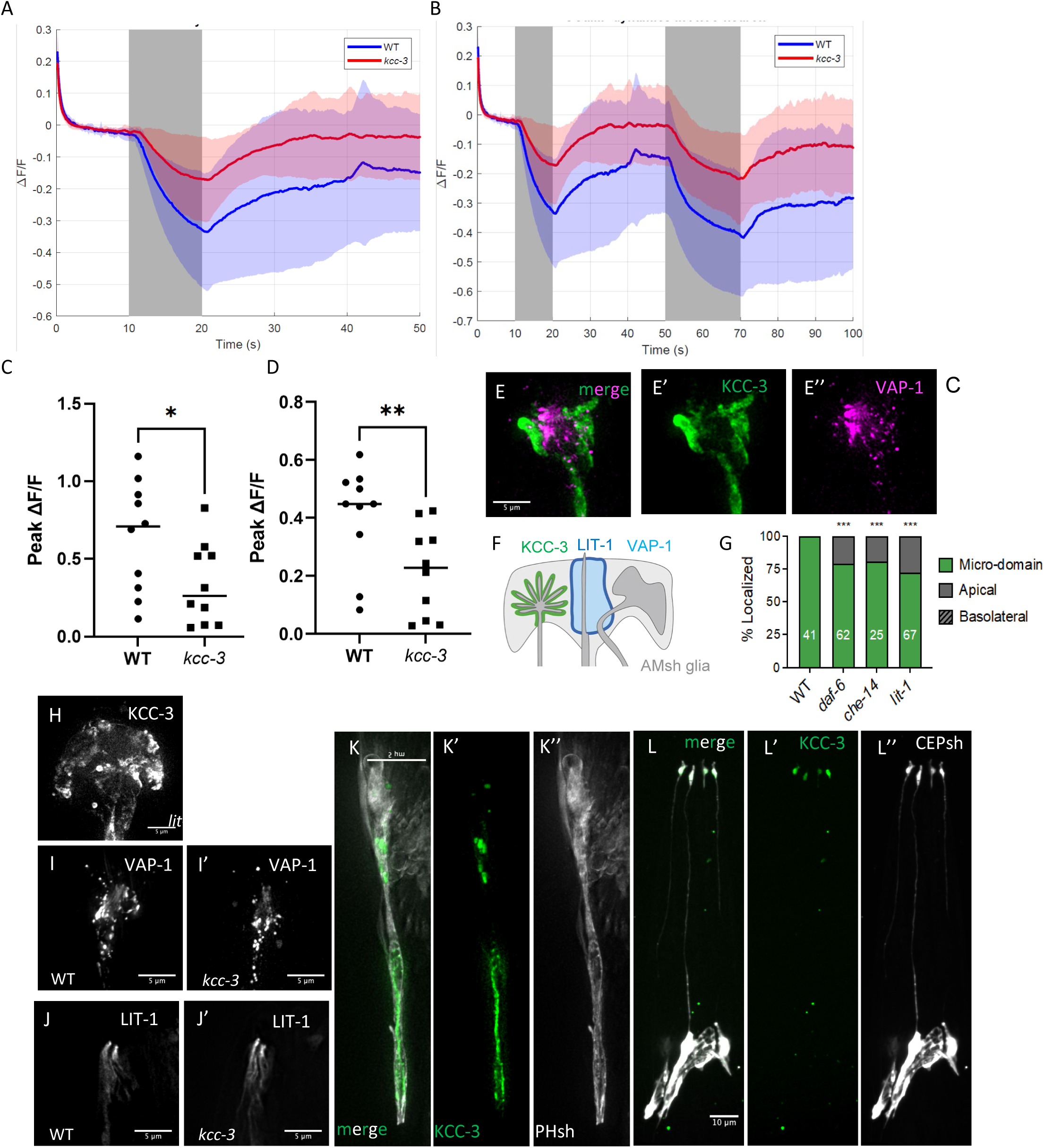
Micro-domains as a general feature of glia. **(A-B)** Calcium transients evoked by addition of 0.01% isoamyl alcohol (IAA) in AWC neuron expressing GCaMP6s upon single (A) or double (B) odor presentations. Solid lines represent the average across 10 different animals in WT (blue) and *kcc-3* (red) background. Shaded areas represent standard deviations. For double odor presentation, odor was presented at 10s and 50s time-points, for 10s and 20s, respectively. N=10 animals **(C-D)** Peak calcium responses when animal presented with 0.01% IAA (p = 0.043; Mann-Whitney) (C). Peak calcium responses when IAA was removed (p = 0.0068; Mann-Whitney) (D). **(E-E’’)** Fluorescence image overlay (E) of KCC-3 (E’) and VAP-1 (E’’). **(F)** Schematic of VAP-1 and LIT-1 localization in AMsh glia, with KCC-3. **(G)** Quantification of KCC-3 localization in *daf-6, che-14, and lit-1* mutants. **(H)** Fluorescent image of KCC-3 in *lit-1* mutants. **(I-I’)** Fluorescent images of VAP-1 in wildtype (I) and *kcc-3* mutant (I’) backgrounds. **(J-J’)** Fluorescent image of LIT-1 in wildtype (J) and *kcc-3* mutant (J’) backgrounds. **(K-L’’)** Fluorescent image overlay (K, L) of KCC-3 (K’, L’) and cytosolic markers (K’’, L’’) in phasmid sheath glia (K-K’’) and CEP sheath glia (L-L’’). *** p < 0.001. All scale bars at 5µm, except L-L’’, which is at 10µm.

### All glial microdomains do not impact distal neurons

KCC-3 regulates AWC-dependent sensory animal behaviors, leading us to wonder if all microdomains can distally regulate other neuron functions. The identity of the AWC molecular microdomain, if any, is unknown, so we asked if channel microdomain cues regulate AFD-NRE. We found that *daf-6* and *vap-1* lesions do not exhibit significant defects in AFD-NRE shape (Figure S7D) and *daf-6* or *che-14* do not impair AFD-mediated thermotaxis behaviors (Perkins et al., 1986). *daf-6* and *che-14* also do not impact sensory behaviors to volatile odorants mediated by wing-neurons (Albert, Brown, & Riddle, 1981; Bargmann, Hartwieg, & Horvitz, 1993). Thus, unlike KCC-3, channel microdomain cues do not modulate distal NREs, suggesting specificity with which these are regulated.

### AMsh glia’s multiple apical microdomains are regulated independently

The results above show that compartmentalized localization of specifically KCC-3 drives cross-modal sensory processing of both AFD-dependent thermotaxis and AWC-dependent chemotaxis behaviors. How is this coordinated within a glial cell? We tested the hypothesis that KCC-3 may do so by altering other glial microdomains.

Previously, it has been shown that the AMsh glia localizes the secreted molecule VAP-1 and membrane-associated LIT-1/NEMO-like kinase at the channel (Oikonomou et al., 2011; Perens & Shaham, 2005). We find that KCC-3 localizes around AFD-NRE. Together, this implies that AMsh glia make at least three molecular microdomains – around channel NREs (VAP-1/LIT-1 positive, KCC-3 negative), AFD-NRE (VAP-1/LIT-1 negative, KCC-3 positive), and wing neurons (VAP-1/LIT-1 negative, KCC-3 negative). To confirm this, we engineered transgenic animals that simultaneously labeled these glial cues and found that KCC-3 indeed localizes to an anterior microdomain distinct from either VAP-1 or LIT-1 (Figure 6E-F). Further, as expected, VAP-1 localizes to an anterior microdomain distinct from AWC (Figure S7E-E’’).

Since the KCC-3 and VAP-1 domains are mutually exclusive, we wondered if loss of either microdomain affected the other. First, we examined if mutations in amphid channel-localized proteins alter KCC-3 localization. We found that mutations in *daf-6, che-14,* and *lit-1,* but not *snx-1,* affect KCC-3 localization, albeit only marginally (Figure 6G-H). In corollary, we also asked if mutations in *kcc-3* reciprocally alter LIT-1 or VAP-1 expression. Again, we did not observe obvious defects (Figure 6I-J’). Thus, while AMsh glial KCC-3 regulates the function of at least the distal AWC neuron, the glia largely maintains its molecular microdomains between AFD and channel neurons independently.

### KCC-3 localizes to apical micro-domains across multiple glia

KCC-3 is broadly expressed in many *C. elegans* glia (Tanis, Bellemer, Moresco, Forbush, & Koelle, 2009), but none of the other glia contact AWC or AFD-NRE. This led us to wonder if all glia restrict KCC-3 localization similarly to AMsh. To test this, we examined KCC-3 localization in two additional sheath glia: the polarized CEPsh at the anterior head of the animal and the PHsh at the posterior tail of the animal. In both cells, KCC-3 localized to presumptive apical regions of the glia, where it contacts cognate NREs (Figure 6K-L’’). Thus, multiple glia localize the K/Cl transporter KCC-3 to discrete apical domains, indicating that its restricted localization in glia likely has broad functional relevance.

## DISCUSSION

Using the discrete localization of glial KCC-3 around a single neuron-contact site (AFD-NRE) as a facile molecular tool, we uncover that a single glial cell has an apical domain maintained by a boundary zone (GAB). Further, it partitions its apical membrane into multiple and distinct molecular microdomains around individual NRE-contact sites. Focusing on one microdomain cue, the K/Cl transporter KCC-3, our genetic and structure-function studies reveal a two-step model for KCC-3 localization. First, it localizes apically, and is then repelled by non-AFD ciliated NREs, rendering it localized to the AFD-NRE (Figure 7). This mechanism is distinct from previously reported regulators of K/Cl family transporters, and KCC-3 localization is required for it to regulate AFD-NRE. Surprisingly, KCC-3 but not all glial microdomain cues, also impact distal neurons. Thus, microdomain localization is important for the glia to compartmentalize cross-modal sensory processing. Finally, we find that different glial microdomains are partially dependent on each other. Thus, this exquisite sub-cellular organization within a glial cell may inform its ability to integrate information across circuits.

**Figure 7:**
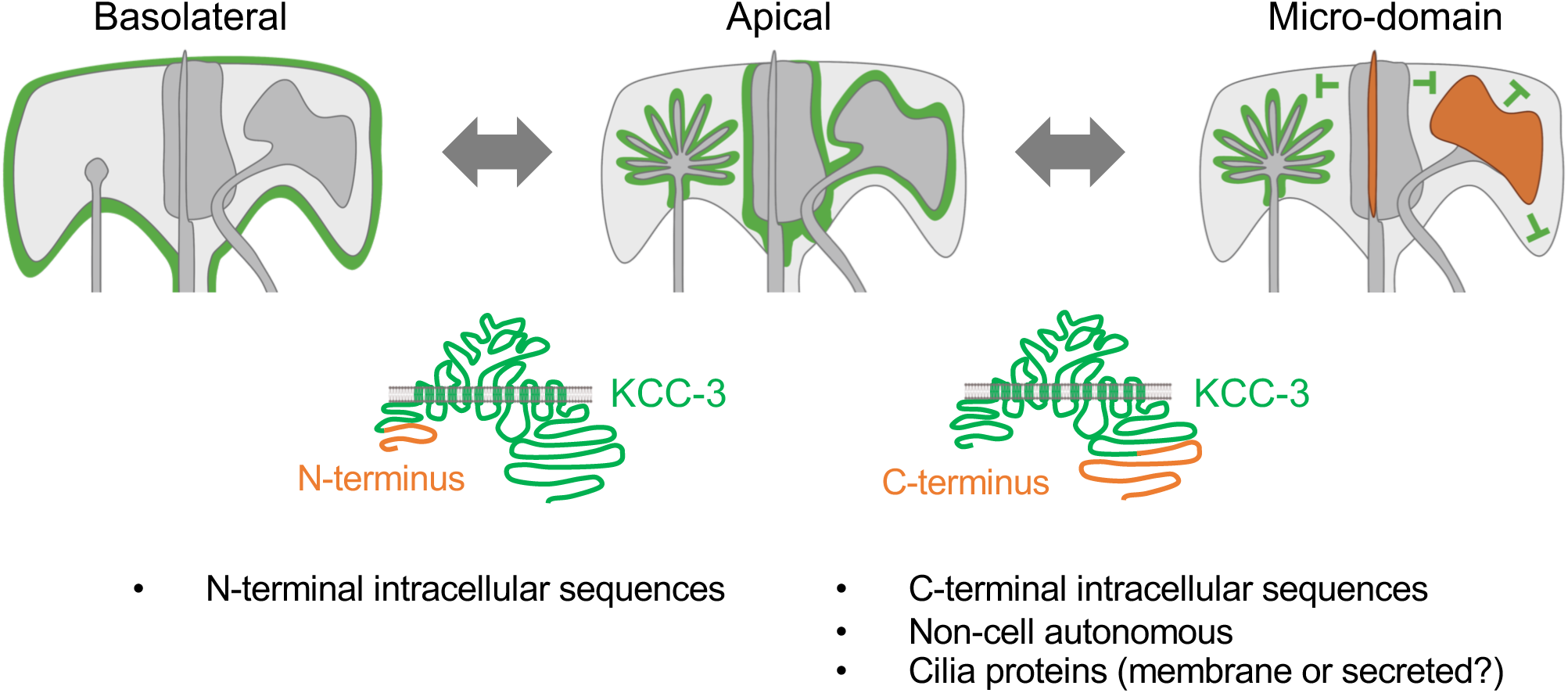
Schematic of KCC-3 localization in AMsh glia. KCC-3 localization is a two-step process. First, N-terminal sequences can guide KCC proteins to basolateral membranes. Second, C-terminal sequences can determine apical vs. micro-domain localization. Cilia also play a role in guiding KCC-3 from broad apical membranes to localized micro-domain membranes.

### Glial apical membranes are molecularly asymmetric at individual neuron contact sites

Our findings track prior studies to show that AMsh exhibit apical-basal polarity (Low et al., 2019). Within even K/Cl co-transporters, we find those that localize either apically (KCC-3) or basolaterally (KCC-1/2). Our studies further identify a 19 AA N-terminal sequence of KCC-2 as necessary and sufficient to drive basolateral localization. In line with prior work on SAX-7, we propose that basolateral targeting is the go/nogo gate that establishes apical-basal polarity for AMsh glia. Sequence overlay does not detect any obvious basolateral-driving motifs shared between SAX-7 and KCC-3, suggesting that multiple motifs may be at play.

The GAB zone boundary in AMsh glia overlays perfectly for all apically restricted molecules (PH-PLCδ, KCC-3, SAX-7) and is not bound by tight junction proteins AJM-1 and DLG-1. This glial cell-biology contrasts with that of epithelial apical-basal polarity and is conceptually analogous to Axon Initial Segments (AIS) in neurons (Leterrier, 2018). We therefore propose that the AMsh glial GAB is a sorting center like the neuronal AIS that delimits diffusion of membrane proteins across different polarized cell domains, with impact on overlay of glia and NRE polarity. How this zone develops or maintains will be interesting to dissect.

### Neuronal cilia signals regulate localization of glial regulatory cues

Most mammalian cells have a primary, non-motile cilia. In neurons and glia, their presence and functions are only recently being appreciated (Green & Mykytyn, 2014; Ki, Jeong, & Lee, 2021; Sengupta, 2017). We find here that non-AFD ciliated NREs regulate localization of a glial transporter at contacting glial membranes, through a signal transported by IFTA/B complex. While further work is needed to identify the molecular identity of this cue, our data hint that this may be independent of proximal extra-cellular vesicle release (Razzauti & Laurent, 2021). To our knowledge, while it has been demonstrated that glia track neuron activity (Agarwal et al., 2017; Duan et al., 2020; Wang, D’Urso, & Bianchi, 2012; Yu et al., 2018), a role for neuronal cilia in guiding glial properties and molecules has not yet been demonstrated.

### Glial microdomain localization of K/Cl transporters

KCC-3 is a SLC12A6/K-Cl electroneutral transporter broadly implicated in neurological diseases including autism, epilepsy, and schizophrenia (Boettger et al., 2003; Delpire & Kahle, 2017; Garneau et al., 2017; Shekarabi et al., 2012). We previously showed that KCC-3 is a glial regulator of neuron shape and functions (Singhvi et al., 2016). Here, we report that glial KCC-3 localizes to a microdomain around only AFD-NRE contact sites. Intriguingly, glia across species localize KCC-3 to molecular microdomains. In rodents, Schwann cell peripheral glia localize KCC-3 to apical microvilli around nodes (Sun, Lin, Tzeng, Delpire, & Shen, 2010). In mammals, inner ear Deiter cells (glia-like support cells) localize KCC-3 to basal poles of hair cells (Boettger et al., 2003; Ray & Singhvi, 2021). This is likely a specific regulation of KCC-3 proteins within glia, as we find that KCC-2 does not exhibit this localization when mis-expressed in glia (Figure 3A-D). Further, this exquisitely specific localization is independent of the canonical kinase regulators of K/Cl biology, WNK/SPAK/OSR kinases, which were identified primarily in KCC-2 studies. Thus, how glia regulate KCC-3 is mechanistically distinct from how other cell-types regulate KCC-1/2, underlining the importance of validating gene functions in cell-specific contexts.

### Glial microdomains and cross-modal information processing

We find that AMsh glia create distinct molecular microdomains of regulatory cues at different neuron/NRE contact sites. AMsh glia also produce Ca^2+^ transients in response to different sensory modalities. Thus, AMsh glia present a powerful experimental platform to overlay molecular and functional microdomain activities with single glia-neuron resolution *in vivo*.

We also find that one microdomain-cue, KCC-3, can regulate sensory processing in distal NREs, suggesting that its exquisite sub-cellular localization influences the glia’s ability to compartmentalize regulation of different contacting neurons. As this is not a general property of all microdomain cues, and because we find that microdomains are largely regulated independently, this raises the notion of specificity – how and why do glia regulate each cue differently? And what does KCC-3’s role in regulating both AWC and AFD imply for the animal’s ability to integrate information between these sensory modalities?

Finally, glia in both peripheral and central nervous systems interact with multiple neurons, and mammalian astrocyte glia exhibit distinct microdomain patterns of intra-cellular Ca^2+^ transients to different neuron activities (Agarwal et al., 2017; Khakh & Sofroniew, 2015). Whether functional Ca^2+^ and molecular microdomains overlay causally awaits inquiry, but already leads us to speculate that their overlay positions glia as integrators of information processing across neural circuits.

## ACKNOWLEDGEMENTS

We thank the Singhvi lab and Jihong Bai for discussions and comments on the manuscript; lab members Cecilia Martin, James Bent, Alex Neitz, and Olivia Okamoto for gift of reagents. We thank Shai Shaham, Max Heiman and Jihong Bai for sharing reagents, and Bai lab for generous support on the calcium imaging studies. This work was funded by Simons Foundation/SFARI grant (488574), Esther A. & Joseph Klingenstein Fund and the Simons Foundation Award in Neuroscience (488574) and NIH/NINDS funding (NS114222) to AS. This work was performed while AS was a Glenn Foundation for Medical Research and AFAR Junior Faculty Grant Awardee. AS sincerely thanks philanthropic supporters to her laboratory including Stephanus and Van Sloun Foundations. Some work was performed at the Fred Hutch Shared Resources Core Facilities. We sincerely apologize if we missed citing works due to our oversight or space considerations.

## AUTHOR CONTRIBUTIONS

SR and AS designed all studies, analyzed data and co-wrote the manuscript. SR performed all experiments and was assisted by RSM and AK in construction of some strains and plasmids. PG performed the functional Ca2+ imaging and analyzed the data with SR and AS.

## Notes

### Competing Interest Statement

The authors have declared no competing interest.

