## Supplemental Materials and Figures for "Neuron cilia constrain glial regulators to microdomains around distal neurons"

This PDF file includes:

Materials and Methods

Figures. S1 to S8

Supplementary Materials Reference List

### EXPERIMENTAL PROCEDURES

#### *C. elegans* methods

*C. elegans* were cultured as previously described (Brenner, 1974; Stiernagle, 2006). Bristol N2 strain was used as wild type. Animals were raised at 20ºC (unless noted) for at least one week without starvation. L4 larval animals were picked to fresh plates and assayed 24 hours later, unless otherwise noted. Germ-line transformations by micro-injection to generate unstable extra-chromosomal array transgenes were carried out using standard protocols (Mello, Kramer, Stinchcomb, & Ambros, 1991). Integration of extra-chromosomal arrays was performed using UV. All transgenic arrays were generated with 30 ng/µl $P_{mig\text{-}24}$:Venus, 15 ng/uL $P_{unc\text{-}122}$:GFP or 20 ng/µL $P_{unc\text{-}122}$:RFP as co-injection markers (Abraham, Lu, & Shaham, 2007; Armenti, Lohmer, Sherwood, & Nance, 2014; Miyabayashi, Palfreyman, Sluder, Slack, & Sengupta, 1999). Information on all strains and reagents is available by request.

#### Strains

Strains were sourced from (a) the CGC, funded by NIH Office of Research Infrastructure Programs (P40 OD010440), (b) the International C. elegans Gene Knockout Consortium (C. elegans Gene Knockout Facility at the Oklahoma Medical Research Foundation, funded by the National Institutes of Health; and the C. elegans Reverse Genetics Core Facility at the University of British Columbia, funded by the Canadian Institute for Health Research, Genome Canada, Genome BC, the Michael Smith Foundation, and the National Institutes of Health) and (c) National BioResource Project (NBRP), Japan.

#### Plasmids

$P_{AMsh}$:KCC-3:worm-mScarlet and $P_{AMsh}$:CFP:SL2:KCC-3:worm-mScarlet: Codon-optimized worm mScarlet was synthesized by Genewiz with flanking AgeI/EcoRI digestion sites and cloned into pAB44 using AgeI/EcoRI to make pSR1/2 (pF53:worm-mScarlet). *mScarlet* and the *unc-54* 3’UTR was cloned into pAS272 ($P_{AMsh}$:KCC-3:mcherry) using Age1/Apa1 from pSR1/2 to make pSR5/6 ($P_{AMsh}$:KCC-3:mScarlet [out of frame]). Site-directed mutagenesis was used to add TATG in front of mScarlet in pSR5/6 to make pSR7/8 ($P_{AMsh}$ :KCC-3:mScarlet [in frame]).

CFP was PCR amplified from pIL43 (gift from Max Heiman) with flanking FseI/XbaI sites and inserted into pAS548 (pSM:SL2:mCherry) to make pSR9/10 (pSM:CFP:SL2:mCherry). CFP-SL2 was then PCR amplified from pSR9/10 and inserted into pSR7 ($P_{AMsh}$:KCC-3:mScarlet) using FseI/AscI to make pSR11/12 ($P_{AMsh}$ :CFP:SL2:KCC-3:mScarlet).

KCC-1/KCC-2: cDNA for *C. elegans* KCC-1 and KCC-2 were PCR amplified in one (KCC-2) or two (KCC-1) segments from a mixed stage cDNA library and cloned into pSR7 using BamH1/EcoRI (KCC-1) or BamH1/SalI (KCC-2) to generate pSR17 ($P_{AMsh}$:KCC-1:mScarlet) and pSR61/62 ($P_{AMsh}$:KCC-2:mScarlet).

DYF-11 rescue constructs: All promoters were PCR amplified from gDNA of mixed stage animals and inserted into pAN1 ($P_{DYF-11}$:DYF-11:GFP) using either SphI/SalI or Gibson assembly. Promoters cloned: $P_{GPA-3}$ (5.4 kb upstream start codon), $P_{TAX-4}$ (3.1 kb upstream start codon), $P_{FLP-19}$ (3.6 kb upstream start codon), $P_{ODR-1}$ (2.4 kb upstream start codon), $P_{R102.2}$ (602 bp upstream start codon), $P_{CEH-36}$ (3kb upstream start codon), and $P_{ODR-4}$ (2.3 kb upstream start codon).

**KCC-2/KCC-3 Chimera Generation**

We used a PCR fusion based approach to create KCC-2/KCC-3 chimera proteins (Hobert, 2018). Briefly, 2 or more segments of KCC-2 or KCC-3 were PCR amplified with nested primers. The 3' primer for every segment but the last included a 24bp overhang to the following segment. In the first PCR, all KCC segments are amplified independently. In the second PCR, nested primers fuse the independent segments to create one contiguous KCC chimeric protein. This product is then inserted into pSR7 using BamH1/Sal1 sites to create the final plasmid.

***dyf-11*** **Rescue Experiments**

All rescue constructs were injected into ASJ583 (*dyf-11(mn392);* dnaIs10 [$P_{AMsh}$:KCC-3:mScarlet]). To blind scoring, both extra-chromosomal array-positive and negative animals were assessed for KCC-3 localization first and then for presence of the rescue construct.

**Statistical Methods**

All data and statistics were graphed and analyzed using Prism9. Proportional data is presented as the proportional sum across multiple days of data collection ± 95% confidence interval. Two-Sided Fisher's Exact tests were performed to compare across genotypes.

**Microscopy, Image Processing and Analysis**

Worms were immobilized with 40mM sodium azide. Images were collected on a Deltavision Elite RoHS wide-field deconvolution system, 40x/1.3 NA oil-immersion or OLY 100×/1.40 NA oil-immersion objective and a DV Elite CMOS Camera. Some images were also captured on a VisiTech iSIM super resolution microscope. Image processing was done in FIJI ImageJ.

**Behavioral Assays**

All chemotaxis behavioral assays as previously described (Bargmann, Hartwieg, & Horvitz, 1993). All assays performed on day 1 adult animals. Briefly, animals are placed at the black dot and allowed to explore for 1hr. 1 μL odorant diluted in ethanol is placed at + and 1 μL ethanol is placed at –. 1 μL of 1M sodium azide is also placed at both + and – points to anesthetize animals. Statistical analysis was performed with unpaired t-test (Graphpad).

**Calcium imaging**

Calcium imaging was conducted in microfluidic chambers (Chronis, Zimmer, & Bargmann, 2007). Imaging was performed at 7 Hz on a Leica DMi8 inverted microscope with a 63x/1.40 NA oil immersion objective and an Andor iXon Life 888 EMCCD camera, using Leica LAS-X software. L4 animals were picked 24 hours prior to imaging and left in 20°C overnight. Animals were immobilized using 1 mM levamisole prior to being loaded into the imaging chamber. Animals were presented with alternating S-Basal buffer and 0.01% isoamyl alcohol diluted in S-basal buffer. Image analysis was performed using ImageJ. ΔF/F was calculated by $(F-F_0)/F_0$, where $F_0$ was designated by the average fluorescent intensity prior to odor presentation (t=1s - t=10s).

**Figure S1. KCC-3 localizes to an apical region specifically around AFD-NRE**

**(A)** Fluorescence image of KCC-3 translational reporter in day 1 adults. **(B-B'')** Fluorescence images as merge (B) and single-channel images of KCC-3 (green, B') and AWB-NRE (red, B''). Non-overlap is denoted by white arrow, site of KCC-3 enrichment around AFD-NRE is noted by magenta asterisk. Scale bar, 5μm.

**Figure S2. KCC-3 localizes to a glial apical microdomain in age-dependent manner**

**(A-C)** Fluorescent images of AMsh membranes tagged with BasoRed (A), ApiGreen (B), and a PH-PLC apical marker (C). **(D-D'')** Fluorescent image overlay (D) of KCC-3 (D') and the junctional DLG-1 (D'') marker. **(E)** Quantification of KCC-3 length expressed as ratio over animal length from nosetip to the first pharyngeal bulb. **** $p < 0.0001$. **(F-G)** Fluorescent images showing apical localization and GAB boundary (white arrow) of KCC-3 in a translational reporter in 3-fold stage embryos (G) and L1 larvae (H).

**Figure S3: Glial KCC-3 localization is regulated by distal non-AFD-NRE cilia**

**(A)** Quantification of KCC-3 localization in Day 1 adults raised at 15°C, 20°C, and 25°C. **(B-B'')** Merge (B) images of $P_{SRTX-1}$ expression (B') in 3-fold embryo (B'', DIC). **(C)** Fluorescent image of KCC-3 expression in Day 1 animals after AFD laser ablation in L1 larvae. **(D)** Fluorescent image of KCC-3 and AFD neuron in Day 1 animals, with AFD on one side ablated. KCC-3 still localizes apically to a micro-domain. **(E)** Schematic of intraflagellar transport (IFT) in *C. elegans*. **(F)** Fluorescent image of KCC-3 localization in *unc-101* mutants. Yellow arrow points to regions that should have AFD enrichment, which is absent. **(G-I)** Fluorescent images of cytosolic CFP in AMsh glia in WT (G), *dyf-11* mutants (H), and *daf-19* mutants (I). **(J)** Schematic of amphid NRE development, with AWB as an example. Ciliary genes, such as *daf-19, dyf-11, and che-11*, interact with identity genes, *ceh-37* in the case of AWB, for the development of the unique NRE shape and function of AWB. **(K)** Quantification of control non-extrachromosomal arrays animals in DYF-11 rescue experiments. All controls are comparable to *dyf-11* mutants.

**Figure S4. Glial KCC-3 microdomain does not require canonical regulators**

**(A)** Quantification of KCC-3 localization in *wnk-1, argk-1,* and *sma-1* loss of function mutants. **(B)** Quantification of KCC-3 localization after RNAi for *wnk-1*, with *pros-1* RNAi as a positive control. **(C-E)** WNK/SPAK phosphorylation sites (C-D) and RFX(V/I) binding domain (E) in human and *C. elegans* KCC-3. These human KCC-3 sites are not conserved in *C. elegans* KCC-3.

**Figure S5. KCC N- and C-terminal swaps**

**(A-B)** Aligned KCC-1, KCC-2, and KCC-3 sequences at the N-terminal (A) and C-terminal regions (B). Dotted lines are where KCC-2 and KCC-3 sequences were swapped to create the chimeras used in the study. Grey box in B denotes region of high sequence dissimilarity at the C-terminal.

**Figure S6: Glial KCC-3 localizes in a two-step process through two protein regions**

**(A)** Description of categories to delineate KCC localization patterns. **(B)** Fluorescent image of KCC chimera localization pattern categorized as "other". **(C-D)** Description (C) and quantification of localization (D) of additional chimeras (10AA swap, 20AA swap), point mutations (LL>AA, ST>AA, TVGE>AVGE, TTS>AAA), and short deletions of KCC-2 and KCC-3. Dotted lines are where KCC-2 and KCC-3 were swapped to create the 10AA and 20AA chimeras, with chimera schematics under the sequence alignment. Point mutations are highlighted, and deletions are denoted by a black bar above or below the sequence that was deleted. The graph (D) is divided into KCC-2 mutations and deletions first and then KCC-3 based chimeras, mutations, and deletions. **(E)** Quantification of KCC chimera localization in WT and *kcc-3* mutant backgrounds. Chimeras tested are denoted under the graph. * $P < 0.05$.

**Figure S7. KCC-3 regulates non-AFD NRE behavior but not shape**

**(A)** Quantification of AWA, AWB, AWC, and ASE collapsed shape in WT and *kcc-3(ok228)* animals. **(B)** Chemotaxis index quantifications for AMsh ablated animals and *ttx-1* mutant animals for odorants and tastant tested in Figure 5. **(C)** Chemotaxis index quantification of *dyf-11* cilia mutants to 1% methyl pyrazine, 1% isoamyl alcohol, and 100mM NaCl. Same day CI values of N2 and *kcc-3(ok228)* mutants included for comparison. **(D)** Quantification of AFD microvilli shape in *daf-6* and *vap-1* mutant animals. **(E-E'')** Fluorescent merge image (E) of AMsh VAP-1 (E') and AWC (E''). These components take up different regions of the AMsh glia.

### REAGENTS

#### A. Mutants

LG1: *tax-2(p691), unc-101(m1), che-1(p678, che-14(ok193)*

LG2: *kcc-3(ok228), daf-19(m86), oig-8(ot818)*

LG3: *lit-1(ns132)*

LG4: *osm-3(p802), gcy-8(ns335)*

LG5: *osm-6(p811), ttx-1(p767), che-11(e1810), che-12(e1812), unc-23(e25), bbs-8(nx77), sma-1(e30)*

LGX: *dyf-11(mn392), odr-7(ky4), ceh-36(ky646), ceh-37(ok642), daf-6(e1377)*

#### B. Integrated transgenes

| Strain | Chromosome | Genotype | Reference |
|---|---|---|---|
| *dnaIs10* | V | $P_{F53F4.13}$*:KCC-3:mScarlet* | This study |
| *dnaIs15* | IV | $P_{F53F4.13}$*:CFP:SL2KCC-3:mScarlet* | This study |
| *nsIs228* | I | $P_{SRTX-1}$*:GFP* | (Singhvi et al., 2016) |
| *ntIs1* | V | $P_{GCY-5}$*:GFP* | (Sarin et al., 2007) |
| *kyIs37* | II | $P_{ODR-10}$*:GFP* | (Sengupta, Chou, & Bargmann, 1996) |
| *kyIs140* | I | $P_{STR-2}$*:GFP* | (Troemel, Sagasti, & Bargmann, 1999) |
| *kyIs104* | X | $P_{STR-1}$*:GFP* | (Troemel, Kimmel, & Bargmann, 1997) |
| *oyIs87* | III | $P_{GPA-4delta6}$*:myr:GFP* | (Maurya & Sengupta, 2021) |

| | | | |
|---|---|---|---|
| *dnaIs19* | | $P_{F53F4.13}$*:SAX-7deltacyt:sfGFP* | (Martin, Bent, & Singhvi, 2022) |
| *hmnIs30* | | $P_{GCY-5}$*:AJM-1:YFP* | Gift from Max Heiman |
| *xnIs17* | | $P_{DLG-1}$*:DLG-1:GFP* | (Firestein & Rongo, 2001) |

**C. Extra-chromosomal arrays**

| **Strain** | **Genotype** | **Reference** |
|---|---|---|
| nsEx2606 | $P_{TO2B11.3}$*:GFP:LIT-1* | (Oikonomou et al., 2011) |
| nsEx4131 | $P_{VAP-1}$*:VAP-1:sfGFP* | Gift from Shai Shaham |
| nsEx5658 | $P_{KCC-3}$*:KCC-3:GFP* (recombineered fosmid) | (Singhvi et al., 2016) |
| nsEx4394 | $P_{SRTX-1B}$*:DYF-11:GFP* | |

**C. Extrachromosomal transgenes and plasmids generated in this study**

* denotes plasmids made outside of the study, which were injected to make stable extra-chromosomal arrays

| **Extra-chromosomal array (dnaEx) number** | **Plasmid** | **Genotype** |
|---|---|---|
| 26, 312, 313 | pSR7 | $P_{F53F4.13}$*:KCC-3:mScarlet* |
| 52, 53, 54, 55, 56, 68 | pSR11 | $P_{F53F4.13}$*:CFP:SL2KCC-3:mScarlet* |
| 222, 223, 224 | pCM11* | $P_{F53F4.13}$*:SAX-7:mApple* |

| | | |
|---|---|---|
| 256, 258, 320 | pCM11* + Recombineered fosmid | *$P_{F53F4.13}$:SAX-7:mApple; $P_{KCC-3}$:KCC-3:GFP* |
| 515, 543, 544 | pOO1 | *$P_{F53F4.13}$:PH-PLCdelta:GFP* |
| 77, 78, 79 | pJF48* | *$P_{SRTX-1}$:EGL-1* |
| 220, 221, 228 | pAN1 | *$P_{DYF-11}$:DYF-11:GFP* |
| 333, 337, 338 | pSR57 | *$P_{GPA-3}$:DYF-11:GFP* |
| 317, 318, 319 | pSR53 | *$P_{TAX-4}$:DYF-11:GFP* |
| 204, 207, 208 | pSR35 | *$P_{ODR-1}$:DYF-11:GFP* |
| 334, 335, 336 | pSR35 + pSR56 | *$P_{ODR-1}$:DYF-11:GFP; $P_{FLP-19}$:DYF-11:GFP* |
| 450, 453, 488 | pSR85 | *$P_{R102.2}$:DYF-11:GFP* |
| 326, 327, 328 | pSR52 | *$P_{ODR-4}$:DYF-11:GFP* |
| | | *$P_{CEH-36}$:DYF-11:GFP* |
| 84 | pSR17 | *$P_{F53F4.13}$:KCC-1a:mScarlet* |
| 75, 76 | pSR15 | *$P_{F53F4.13}$:KCC-2a:mScarlet* |
| 153 | pSR24 | *$P_{F53F4.13}$:Chimera A:mScarlet* |
| 188, 189, 193 | pSR27 | *$P_{F53F4.13}$:Chimera B:mScarlet* |
| 194, 206 | pSR33/34 | *$P_{F53F4.13}$:Chimera C:mScarlet* |
| 314, 315, 316 | pSR43 | *$P_{F53F4.13}$:Chimera D:mScarlet* |
| 325, 351, 352 | pSR45 | *$P_{F53F4.13}$:Chimera E:mScarlet* |
| 167, 454, 462, 484 | pSR25 | *$P_{F53F4.13}$:Chimera F:mScarlet* |
| 410, 411, 417 | pSR83 | *$P_{F53F4.13}$:Chimera G:mScarlet* |
| 445, 446, 447 | pSR87 | *$P_{F53F4.13}$:Chimera H:mScarlet* |

| | | |
|---|---|---|
| 252, 257, 302 | pSR39 | $P_{F53F4.13}$*:KCC-2 LL>AA:mScarlet* |
| 332, 342, 381 | pSR61 | $P_{F53F4.13}$*:KCC-3 TTS>AAA:mScarlet* |
| 253, 254, 255 | pSR41 | $P_{F53F4.13}$*:KCC-3 ST>AA:mScarlet* |
| 339, 340, 341 | pSR59 | $P_{F53F4.13}$*:KCC-3 TVGE>AVGE:mScarlet* |
| 384, 385, 386 | pSR69 | $P_{F53F4.13}$*:KCC-2 deletion 1:mScarlet* |
| 412, 413, 415 | pSR71 | $P_{F53F4.13}$*:KCC-2 deletion 2:mScarlet* |
| 428, 430 | pSR66 | $P_{F53F4.13}$*:KCC-3 deletion 3:mScarlet* |
| 407, 427, 429 | pSR73 | $P_{F53F4.13}$*:KCC-3 deletion 4:mScarlet* |
| 383, 387, 388 | pSR67 | $P_{F53F4.13}$*:KCC-3 deletion 5:mScarlet* |
| 408, 409, 425, 426 | pSR75 | $P_{F53F4.13}$*:KCC-3 deletion 6:mScarlet* |
| 5, 6 | pSR7 + pCF27* | $P_{F53F4.13}$*:KCC-3:mScarlet;*<br><br>$P_{VAP-1}$*:VAP-1:sfGFP* |
| 514, 517, 524 | pPG5/6 | $P_{HLH-17}$*:KCC-3:mScarlet* |

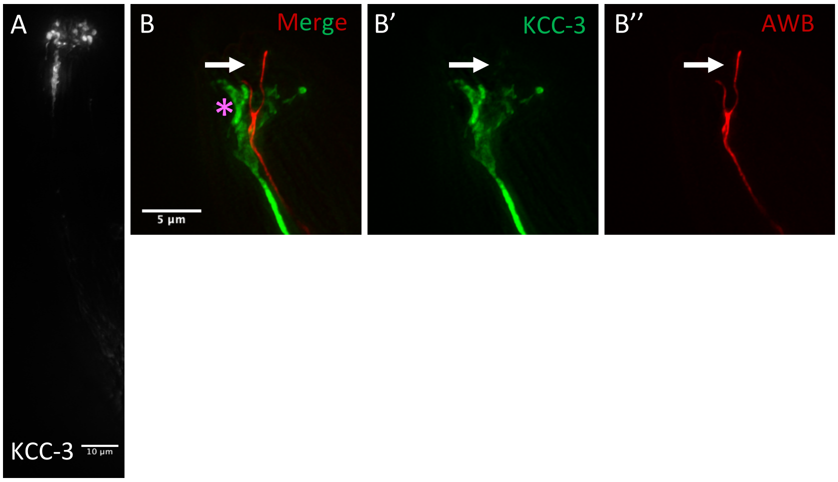
A
KCC-3
10 µm
B
Merge
*
5 µm
B'
KCC-3
B''
AWB

Figure S2

Ray et al

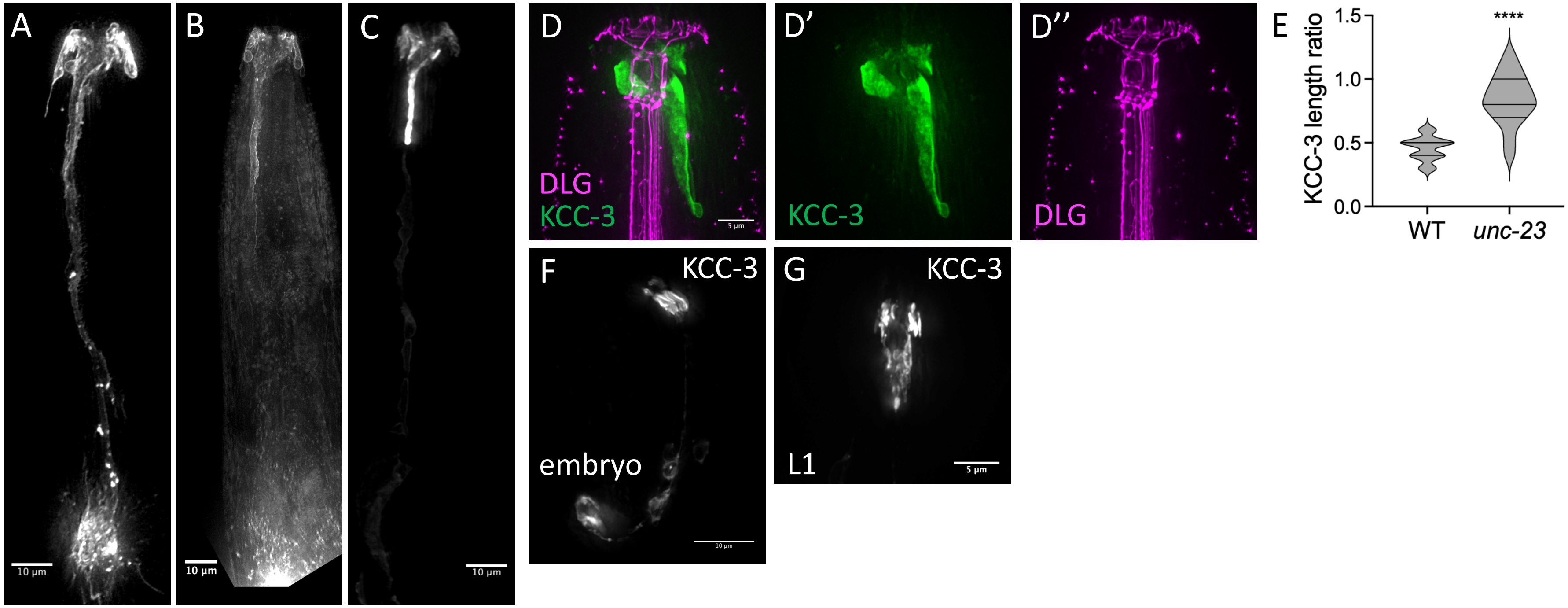

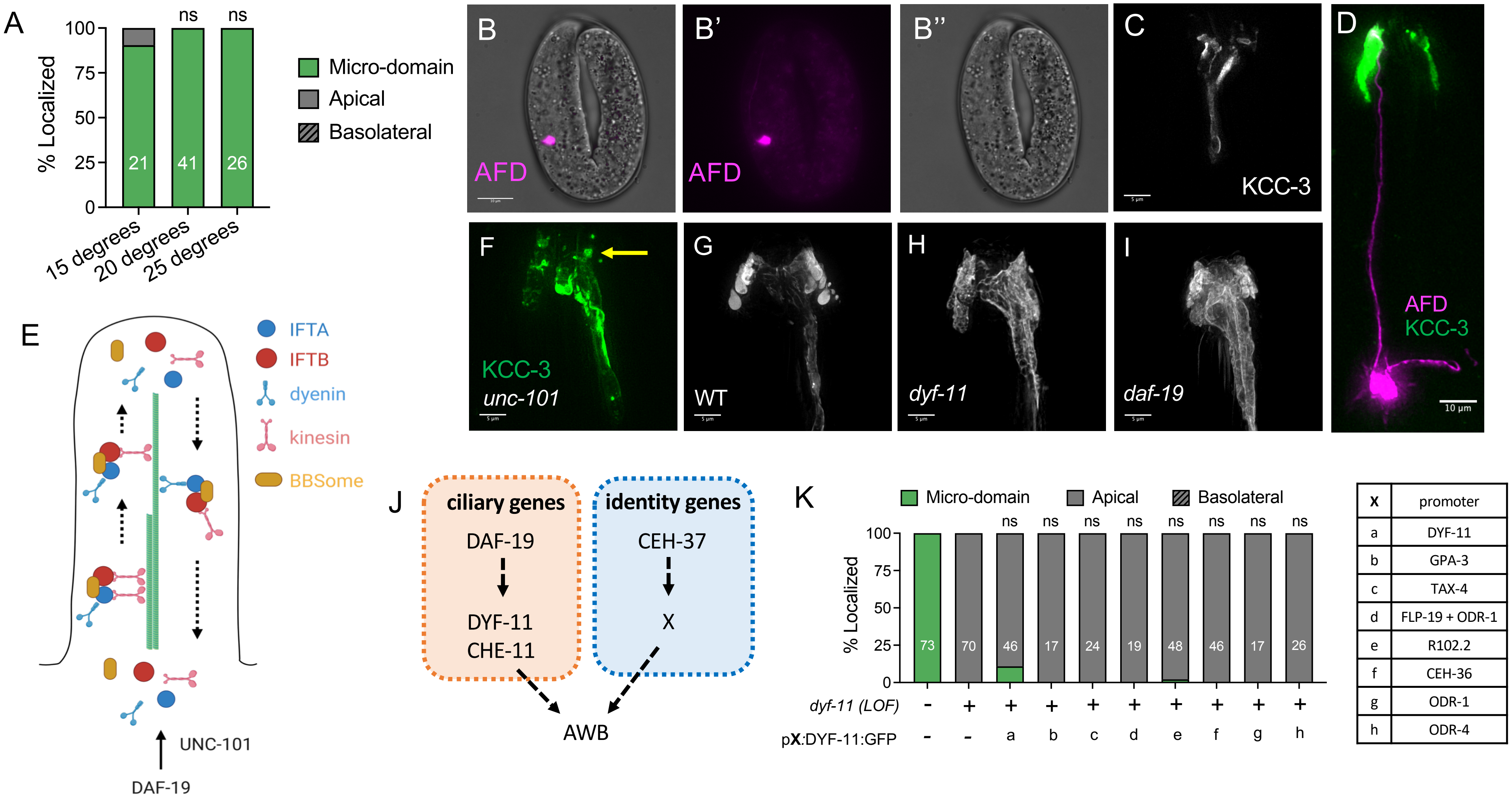


| x | promoter |
|---|---|
| a | DYF-11 |
| b | GPA-3 |
| c | TAX-4 |
| d | FLP-19 + ODR-1 |
| e | R102.2 |
| f | CEH-36 |
| g | ODR-1 |
| h | ODR-4 |

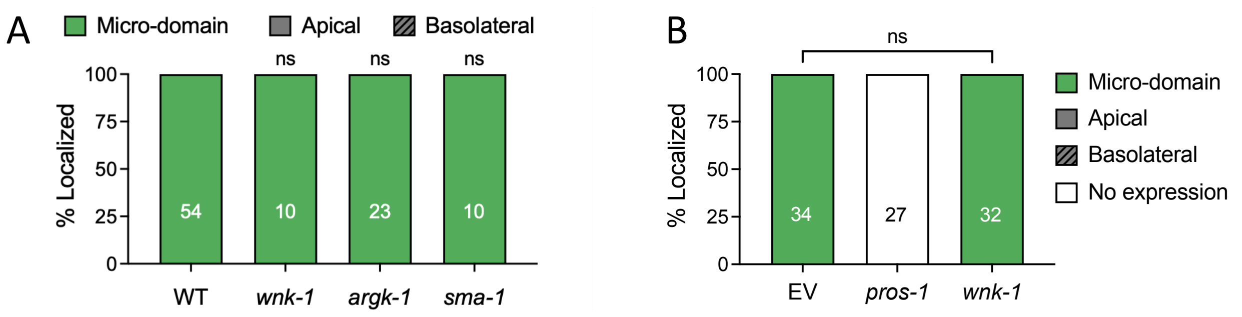


### C Thr991

```
ceKCC3a     IYMLRIDAELFIVDLLDMEVSDEVVEKAA     910
ceKCC3b     IYMLRIDAELFIVDLLDMEVSDEVVEKAA     921
hKCC3a      LYHLRIEAEVEVVEMHDSDISAYTYERTL     992
hKCC3b      LYHLRIEAEVEVVEMHDSDISAYTYERTL     941
hKCC3c      LYHLRIEAEVEVVEMHDSDISAYTYERTL     933
hKCC3d      LYHLRIEAEVEVVEMHDSDISAYTYERTL     983
hKCC3e      LYHLRIEAEVEVVEMHDSDISAYTYERTL     977
            :*  ***:**: :*:: * ::*   . *::
```

### D Thr1048

```
ceKCC3a     ETSFTESLF----DDFY--RSGTP     1088
ceKCC3b     ETSFTESLF----DDFY--RSGTP     999
hKCC3a      TETYQEKVHMTWTKDKYMASRGQK     1061
hKCC3b      TETYQEKVHMTWTKDKYMASRGQK     1010
hKCC3c      TETYQEKVHMTWTKDKYMASRGQK     1002
hKCC3d      TETYQEKVHMTWTKDKYMASRGQK     1052
hKCC3e      TETYQEKVHMTWTKDKYMASRGQK     1046
              :: *.:.    .* *     *
```

### E SPAK/OSR1 RFX(V/I) binding domain

```
ceKCC3a     -----------------------------     0
ceKCC3b     ------------------------MGGFQ     5
hKCC3a      MHPPETTTKMASVRFMVTPTKIDDIPGLS     29
hKCC3b      -----------------------------     0
hKCC3c      -----------------------------     0
hKCC3d      ---------MASVRFMVTPTKIDDIPGLS     20
hKCC3e      MHPPETTTKMASVRFMVTPTKIDDIPGLS     29
```

## A

```
ceKCC1      MGDDSSQDKASAPPSAAAAGAAPAGNPQQQPASAIRNKVDVPPGFGRRVSSVEVVKNAMP   60
ceKCC2      ------------------------------------------------------------   0
ceKCC3a     ------------------------------------------------------------   0

ceKCC1      PTKKASVVIFETPDTPKAERRASSWWRNLVMDDDKTAPVQSDTVISPENAGNGLADH---   117
ceKCC2      --------------MSSE-------------------PRHTFLYVNPNNEAIRLLDIDEG   27
ceKCC3a     --------------MSNARRRFST-----------VTQINTEGL-QAMGKGGGRMET-VG   33
                            .                      ::    .  . .    :
```

Chimera B & C (55AA KCC-3)

```
ceKCC1      HRNISTASVGDRELYAWGTNKDAGGNHNLALF-EEP-----SMPFFSSYLKAHITPGPLE   171
ceKCC2      AERIPL----------------CEQHSPLALYEQDYDTQGQKIGTMLRKLSVYNATE--A   69
ceKCC3a     EDGIPADYKGNRK--------FTTSLGHLALYKEDE-----GIGTQASFISGYTTPGPKE   80
               *                        ***: ::       :      :. : :

ceKCC1      RAQSSSHGGHGAKADLGVLLGVYLPTIQHILGVTMFIRLFWLVGIAGLGQTFLLLFLCCF   231
ceKCC2      TSVEADEKPKAAAAKMGTIMGVFLPCLQNIFGVLFFIRLAWIIGTAGVFQAFFVVLTCVS   129
ceKCC3a     RATS-----EHVKANLGVMLGVYLPTIQHILGVTMFIRLFWVVGMSGVAWTMALLAICCL   135
             : .     . . *.:*.::**:** :*:*:** :**** *::* :*:  :: ::  *
```

predicted N-terminus

Chimera A

## B

```
ceKCC1      LCLLMGYLLKQHKVWRGCKLRVIGIAQE-SDNNVKMQEDLQKYVYQLRIDAKIMIVELAD   982
ceKCC2      LLMLLPFLLRQHKTWKNTTVRLFAIAQM-EDNNVQMKTDLEKFLYHLRIDAAVNVIEMTD   873
ceKCC3a     ILMLIAYLLRQHKVWKGCTLRIFAVSEQDSTKSEDMKAGLQKYIYMLRIDAELFIVDLLD   898
            : :*: :**:***.*:. .:*::.:::  . :. .*: .*:*::* ***** : :::: *
```

Chimera F

C-term high sequence difference (81 AA)

```
ceKCC1      PEISKNAFERTLLMEERTMMMRDLQKVSGGGMSLSLPPAN-APRAPS------PLVTSER   1035
ceKCC2      SDISDYTYERTMKMEERNQYLKNLNKSDRDKDIQNHLEIVTRERKLSRINEEAPAVVPEQ   933
ceKCC3a     MEVSDEVVEKAAEVERKQKEREEMRRSKSGYLNDGFMEDNGKPRQV--------MMRHSD   950
             ::*. . *::  :*.:    .::.: . .    .        *           :  .

ceKCC1      R-ANSKDS--DEGT--PT--ESEETTEKKSTSTDNEQ---ANQETKTKKERMKALDRSKV   1085
ceKCC2      RNLEVVDEEQEDGKSENGSAKIEHKGVRFSDDEDSKEVKVGNGTLERDREERQRKRRYNV   993
ceKCC3a     S-ARSFSP--QPGA--HTSINLDETETSFTES--LFD---DFYRSGTPNEDLEGAMKLNI   1000
               .  .   : *       : :..    : .    :           .*  :   : ::
```

Chimera G

```
ceKCC1      SKMHTAVRLNELLLQHSANSQLILLNLPKPPVHKDQQALDDYVHYLEVMTDKLNRVIFVR   1145
ceKCC2      HKMHTAVKLNELMRQKSSDAQLVFVNLPGPPDADS---DSYYMDFIEALTEGLDRVLLVR   1050
ceKCC3a     HKMNTSVRLNRVIRENSPDSQLILLNLPSPPRNRL-AFNNSYMTYLDVLTEDLPRVLFIG   1059
             **:*:*:**.:: ::* ::**:::*** **        . *: :::.:*: * **:::

ceKCC1      GTGKEVITESS          1156
ceKCC2      GTGAEVVTIYS          1061
ceKCC3a     GSGREVITIDS          1070
            *:* **:*  *
```

A

| Category | Features |
|---|---|
| Basolateral | • GAB absent (entire glial cell)<br>• Outer membrane |
| Apical | • GAB present<br>• Expression in AMsh “wings |
| Micro-domain | • GAB present<br>• Restricted expression |
| Other | • GAB absent (entire glial cell)<br>• Inner membranes |

B

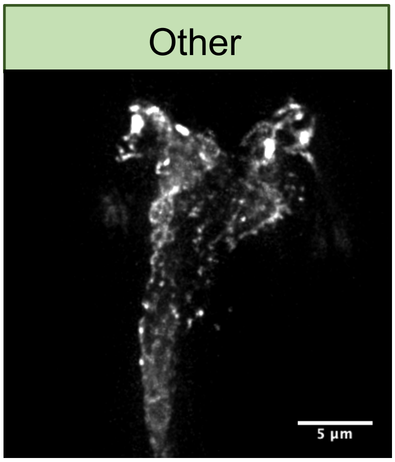


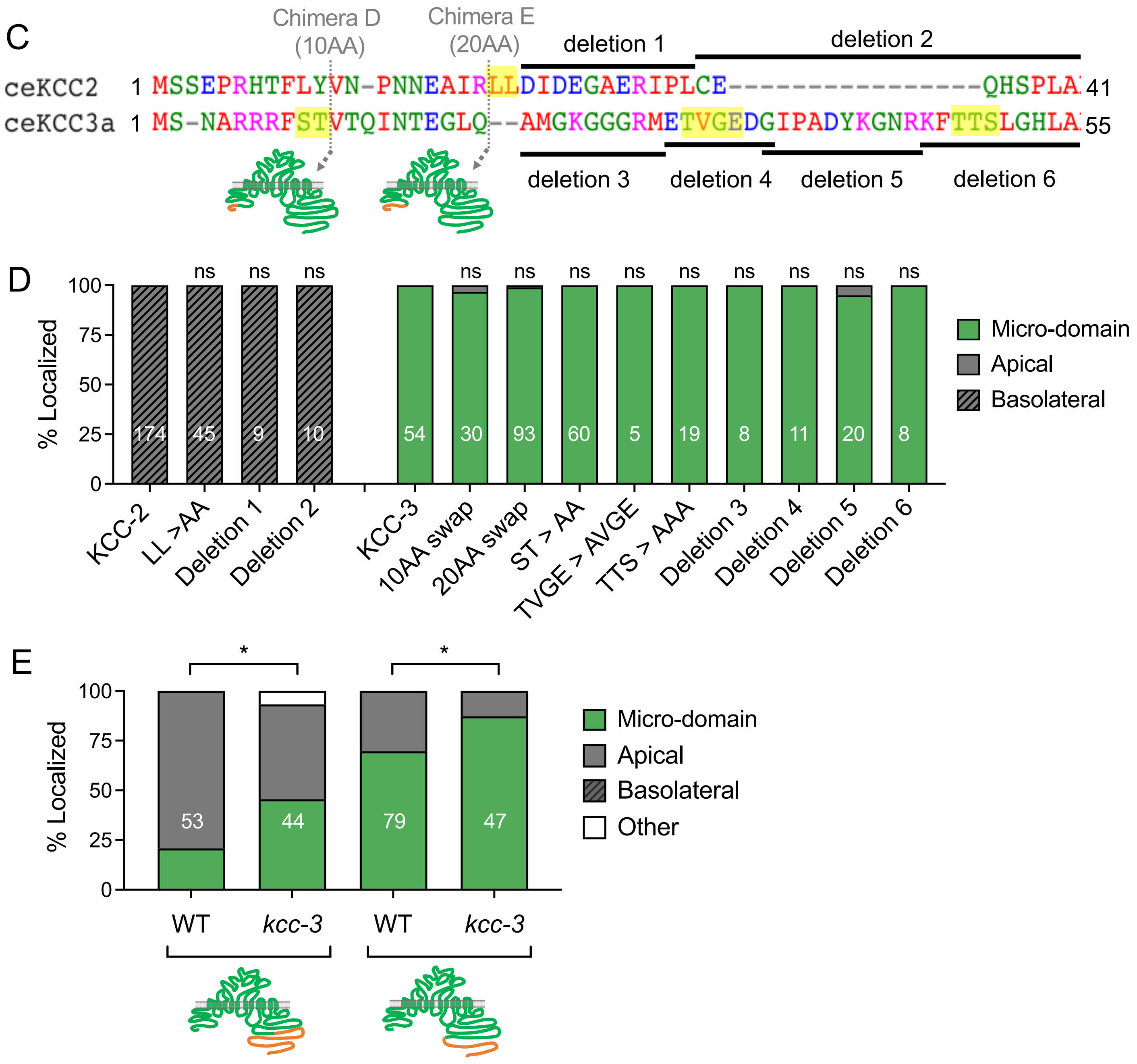

A

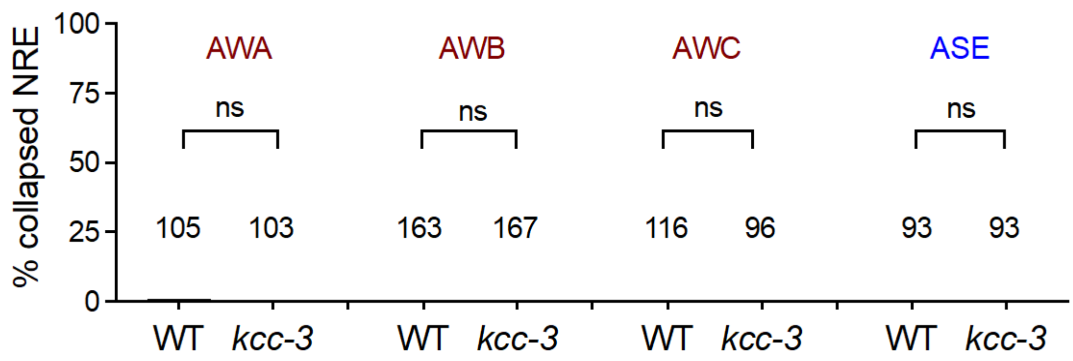


B

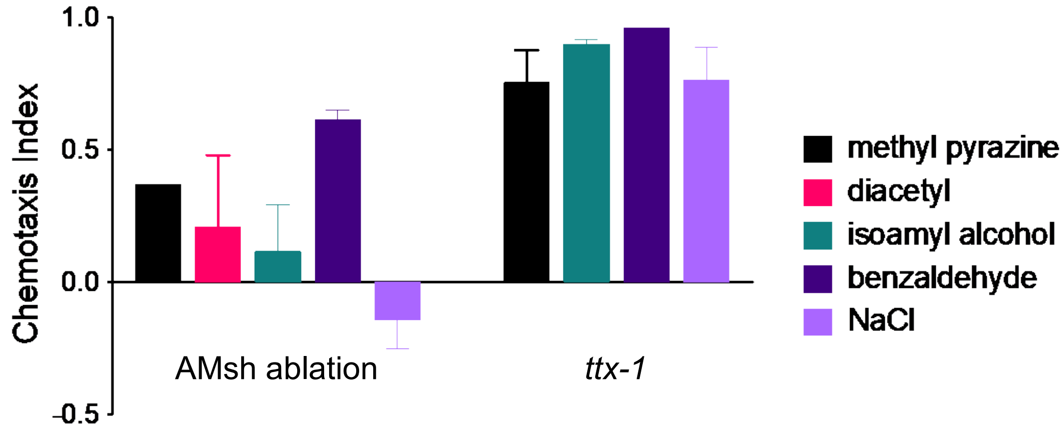


C

| Neuron | Chemoattractant | N2 | *dyf-11* | *kcc-3* |
|---|---|---|---|---|
| AWA | 1% methyl pyrazine | 0.43 | 0.2 | 0.15 |
| AWC | 1% isoamyl alcohol | 0.48 | 0.21 | 0.18 |
| ASE | 100mM NaCl | 0.6 | 0.54 | 0 |

D

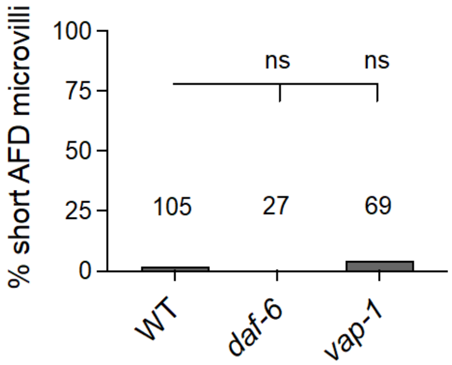


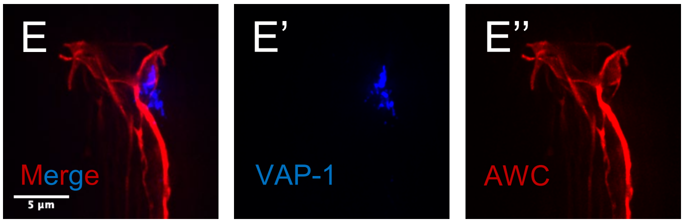
